# Senescence-Induced Immune Remodeling Facilitates Metastatic Adrenal Cancer in a Sex-Dimorphic Manner

**DOI:** 10.1101/2022.04.29.488426

**Authors:** Kate M. Warde, Lihua Liu, Lorenzo J. Smith, Brian K. Lohman, Chris J. Stubben, H. Atakan Ekiz, Julia L. Ammer, Kimber Converso-Baran, Thomas J. Giordano, Gary D. Hammer, Kaitlin J. Basham

## Abstract

Aging is a carcinogen that markedly increases cancer risk, yet we have limited mechanistic understanding of cancer initiation in aged cells. Here, we demonstrate induction of the hallmark aging process cellular senescence, triggered by loss of Wnt inhibitor ZNRF3, remodels the tissue microenvironment and ultimately permits metastatic adrenal cancer. Detailed characterization reveals a striking sexual dimorphism. Males exhibit earlier senescence activation and a greater innate immune response. This results in high myeloid cell accumulation and lower incidence of malignancy. Conversely, females present a dampened immune response and are more prone to metastatic cancer. Senescence-recruited myeloid cells become increasingly depleted with advanced tumor progression, which is recapitulated in patients where a low myeloid signature is associated with worse outcome. Collectively, our study reveals a novel role for myeloid cells in restraining adrenal cancer progression with significant prognostic value, and provides a model for interrogating pleiotropic effects of cellular senescence in cancer.

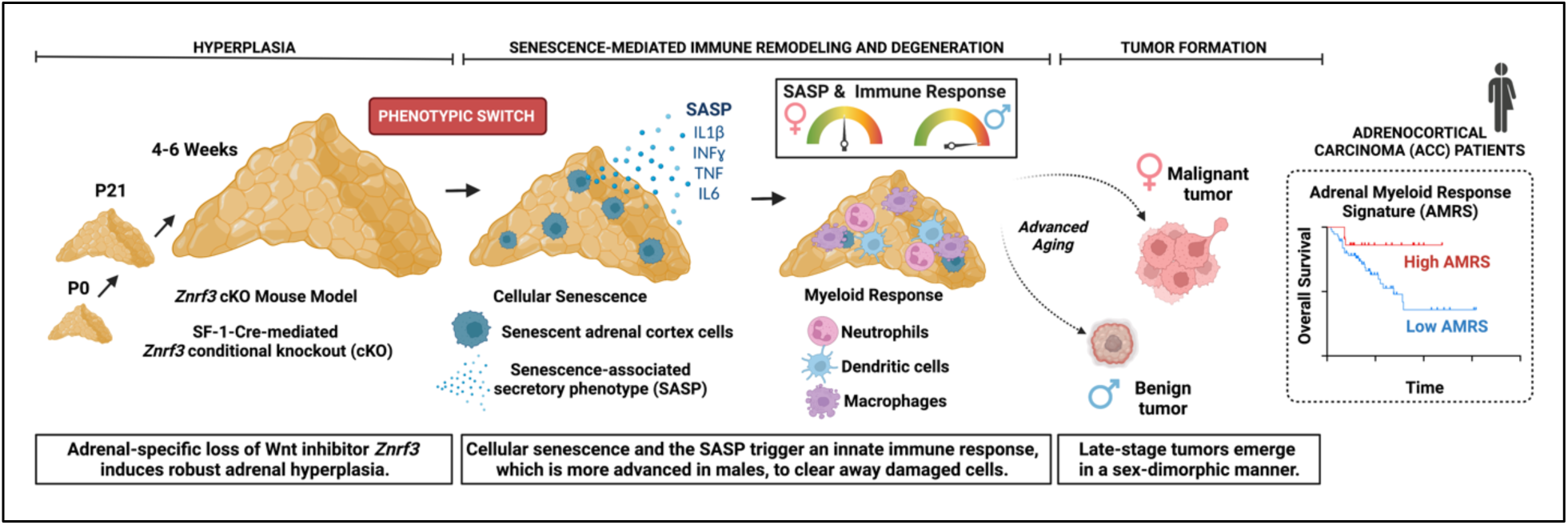
Graphical abstract created with BioRender.com

## Introduction

Aging is the most important risk factor for cancer^1^. Overall incidence rates rise steeply with age, with nearly 90% of newly diagnosed cancer occurring after age 50^2,3^. Age-driven changes in the tissue landscape cooperate with oncogenic mutations within cells to determine overall cancer risk^4^. During aging, the tissue microenvironment evolves based on a multitude of physiological factors, including oxygen and nutrient availability, extracellular matrix (ECM) composition, and immune cell infiltration^5^. These tissue-level changes influence the functional impact of oncogenic mutations that accumulate with age^3^. However, despite aging being a major carcinogen, we have only a limited understanding of how the aged microenvironment cooperates with genetic mutations to influence tumorigenesis.

Adrenocortical carcinoma (ACC) is a highly pertinent model to study the interplay between the aging tumor microenvironment and cell-intrinsic driver mutations. ACC is an aggressive cancer of the adrenal cortex with no effective treatments^6^. Moreover, peak incidence occurs at age 45-55^7^, which corresponds to ∼12-18 months in mice^8^. One pathway that is frequently hyperactivated in ACC is Wnt/β-catenin, a highly conserved signaling cascade that mediates proper cell renewal and cell-to-cell interactions in a wide range of tissues^9^. Unlike many Wnt-driven tumors that are induced by activating mutations in *CTNNB1* (β-catenin) or inactivation of key components of the β-catenin destruction complex (*e.g. APC*), ACC tumors frequently harbor loss-of-function (LOF) alterations in *ZNRF3*^10,11^, an E3 ubiquitin ligase that promotes endocytic turnover of Wnt receptors^12,13^. Consequently, inactivation of *ZNRF3* increases receptor availability and renders cells hypersensitive to Wnt ligands within the microenvironment.

Wnt-driven ACC has been previously modeled in mice through *Ctnnb1* gain-of-function (GOF)^14^ or *Apc* loss-of-function (LOF)^15^. These approaches result in mild hyperplasia with late progression to benign adenomas (>40-weeks) or in some rare cases, carcinomas (17-months, ∼20% penetrance). Combined *Ctnnb1*-GOF and *Tp53*-LOF^16^ shortens tumor latency, but still requires 12-weeks or longer for tumors. These studies hint at an important role for aging in the initiation and progression of adrenal tumors. However, these models are based on less frequent Wnt alterations in ACC, and the mechanism by which aging influences tumorigenesis remained unresolved.

We recently developed a model to specifically ablate *Znrf3* in the adrenal cortex using steroidogenic factor-1(SF-1)-Cre, which mediates high efficiency recombination throughout the adrenal cortex beginning at E9.5^17^. We initially characterized *Znrf3* conditional knockout (cKO) mice between P0 and adrenal maturation at 6-weeks^18^. We observed progressive adrenal cortex hyperplasia that resulted in an average 8.4-fold increase in adrenal weight. Further characterization revealed a gradient of Wnt/β-catenin activity required for normal homeostasis with a specific role for ZNRF3 in limiting expansion of Wnt-low cells. Nevertheless, we found no evidence of neoplastic transformation during the early time frame of these studies.

Here, we investigated the interplay between genetic *Znrf3* inactivation and the aging microenvironment. We hypothesized that SF1-Cre-driven *Znrf3* cKO mice would progress from adrenal hyperplasia to carcinoma with age. Unexpectedly, however, we found that *Znrf3* cKO adrenals steadily regress over time after the initial hyperplastic phase. We demonstrate this phenotypic switch from hyperplasia to regression is driven by activation of cellular senescence and a subsequent senescence-associated secretory phenotype (SASP). We characterize diverse immune cell infiltrates that remodel the tissue microenvironment following senescence activation, which exhibit a highly sex-dimorphic response that is enhanced in males. However, following prolonged inflammation, a large proportion of mice ultimately develop adrenal tumors. Males predominately form benign lesions, while females are significantly more prone to developing metastatic adrenal tumors, mirroring the higher incidence of ACC in women^7^. Myeloid-derived immune cells are excluded from the tissue microenvironment as tumors progress, which is recapitulated in human ACC patients where a low myeloid signature is associated with worse outcome. Taken together, our results suggest that *Znrf3* loss is permissive for ACC tumor progression with advanced age. More broadly, our work suggests a mechanism by which extrinsic factors including sex and evolution of the immune microenvironment can cooperate with genetic lesions to promote age-induced malignant transformation.

## Results

### Ultrasound imaging provides a reliable measure of adrenal size in real-time

Given significant adrenal hyperplasia we previously observed at 6-weeks, we hypothesized that *Znrf3* cKOs would progress to form adrenal tumors with increased age. To test this hypothesis, we sought to monitor adrenal growth in real-time. We employed ultrasound imaging as a non-invasive technique to quantify adrenal size. We first validated this approach using a mixed cohort of mice across a range of ages with varying adrenal sizes. We measured adrenal area and volume by ultrasound, then euthanized mice 24-hours post-imaging to obtain adrenal weight (Extended Data Fig. 1a). We found that adrenal weight was significantly correlated with both ultrasound area (R^2^=0.90, **p=1.08×10^-3^) (Extended Data Fig. 1b) and volume (R^2^=0.89, **p=1.35×10^-3^) (Extended Data Fig. 1c). These results demonstrate that ultrasound imaging provides an accurate assessment of adrenal size in real-time.

### Following initial hyperplasia, *Znrf3*-deficient adrenals regress over time

To monitor for potential tumors, we performed weekly adrenal ultrasound imaging in *Znrf3* cKOs. We began imaging females at 4-weeks and observed an expected increase in adrenal size by 6-weeks (Fig. 1a-b). These results were consistent with our prior studies demonstrating that *Znrf3* loss in the adrenal cortex causes progressive hyperplasia between P0 and 6-weeks^18^. However, rather than continued growth, we unexpectedly observed a gradual decline in adrenal volume after 6-weeks (Fig. 1a-b).

**Figure 1.**
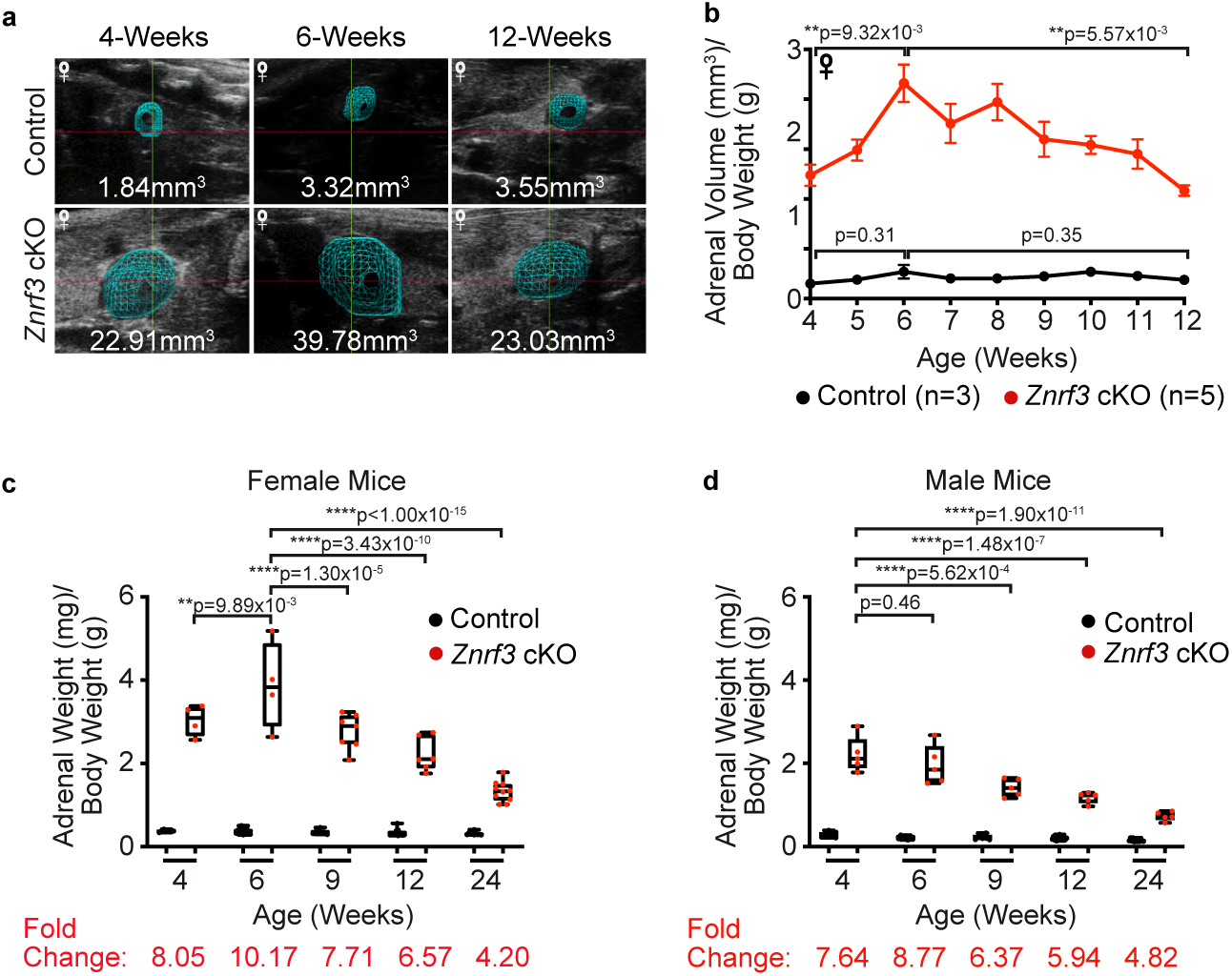
Following an initial phase of significant hyperplasia, *Znrf3*-deficient adrenal glands regress over time. (a) Ultrasound imaging provides a non-invasive method for measuring adrenal size in real-time. Representative 3D ultrasound images from a female control and *Znrf3* cKO mouse are shown at 4-weeks (initiation of study), 6-weeks (maximum volume), and 12-weeks (end of study) of age. (b) Weekly ultrasound monitoring in female control and *Znrf3* cKO mice identifies a phenotypic switch from hyperplasia to regression at 6-weeks of age. Statistical analysis was performed using one-way ANOVA followed by Tukey’s *post hoc* test. Error bars represent mean ± s.e.m. (c) Adrenal weight measurements over time confirm significant adrenal regression in female *Znrf3* cKOs beginning after 6-weeks. (d) In males, adrenal weight peaks earlier than females, and progressively declines with increased age. Each dot represents an individual animal. Box and whisker plots represent mean with variance across quartiles. Statistical analysis was performed using two-way ANOVA followed by Tukey’s multiple comparison test.

To follow-up on these observations, we collected cohorts of male and female mice at 4-, 6-, 9-, 12-, and 24-weeks of age. Consistent with ultrasound measurements, we initially observed increased adrenal weight in female *Znrf3* cKOs between 4- and 6-weeks, followed by a progressive decline in adrenal weight over time (Fig. 1c). In males, adrenal weight did not continue to increase after 4-weeks, and similar to females, gradually declined with increasing age (Fig. 1d). Taken together, these results indicate that *Znrf3*-deficient adrenals undergo a distinct phenotypic switch from hyperplasia to regression.

### Initiation of adrenal regression is marked by reduced proliferation

We hypothesized that declining adrenal size in *Znrf3* cKOs was caused by enhanced cell death. To test this, we measured apoptosis, which is the dominant form of cell death attributed to homeostatic turnover of the normal adrenal cortex^19^. Using immunohistochemistry (IHC) for cleaved caspase 3 (CC3), we found CC3-positive cells primarily localized at the cortical-medullary boundary in controls, as expected (Fig. 2a-b). In *Znrf3* cKOs, apoptotic cells were scattered throughout the cortex, consistent with fragmentation of the medullary compartment in *Znrf3* cKOs as previously described^18^. However, there was no increase in the proportion of CC3-positive cells in *Znrf3* cKOs compared to controls (Fig. 2a-b). We concluded the age-dependent decrease in adrenal size in *Znrf3* cKOs was not caused by increased apoptotic cell death.

**Figure 2.**
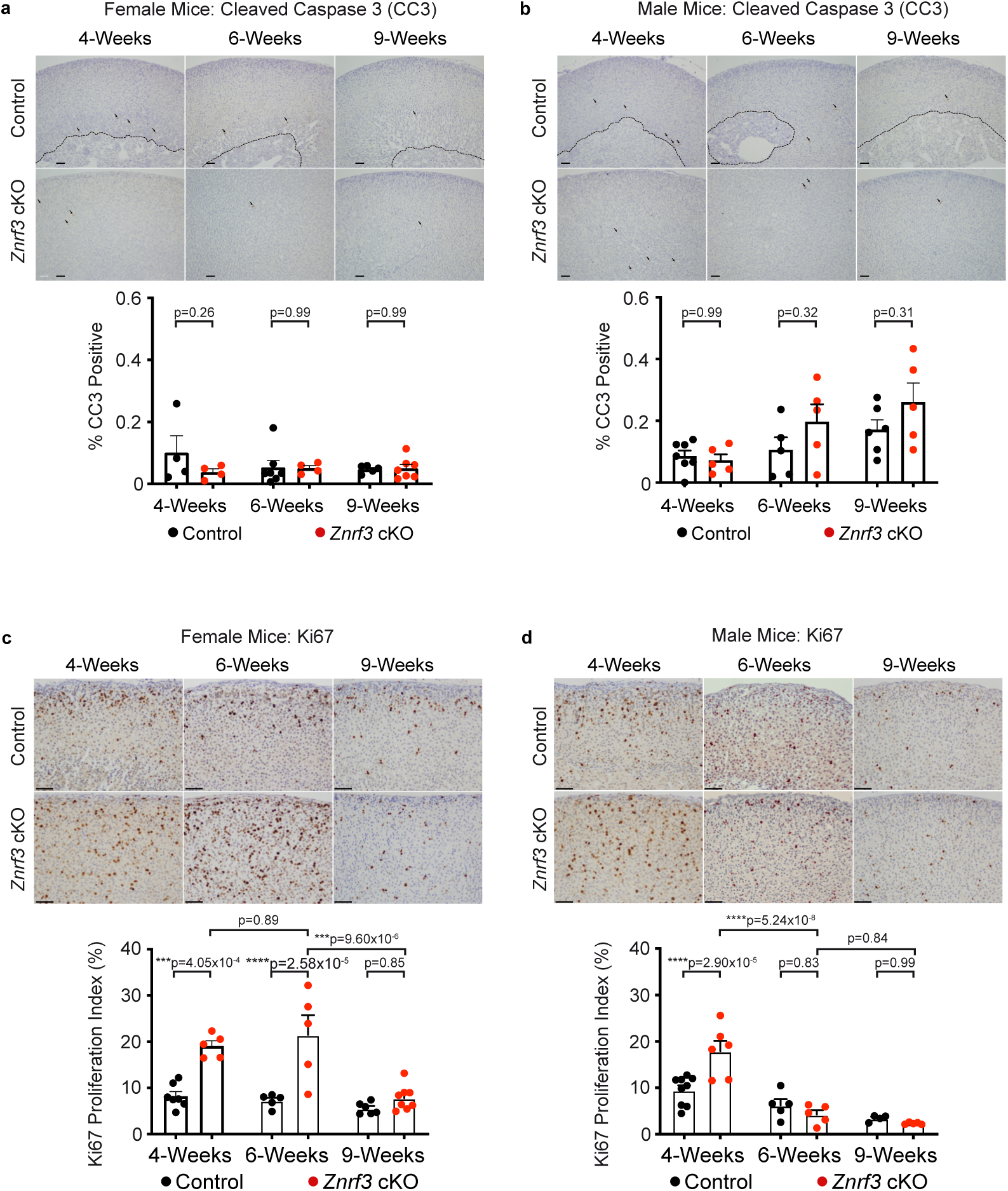
Adrenal gland regression is associated with a significant reduction in proliferation. (a-b) Apoptotic cell death as measured by cleaved caspase 3 (CC3) is not significantly increased in female or male *Znrf3* cKO adrenals compared to controls during the initial phase of tissue regression. Quantification of CC3-positive cells was performed using QuPath digital image analysis based on the number of positive cells and normalized to total adrenal cortex nuclei. Arrows indicate CC3-positive cells. Dashed line indicates histological boundary between adrenal cortex and medulla. (c) Proliferation as measured by Ki67 is significantly increased in 4- and 6-week female *Znrf3* cKO adrenals compared to controls. At the onset of adrenal regression (9-weeks), proliferation is significantly reduced in *Znrf3*-deficient adrenals. (d) Male *Znrf3* cKOs similarly exhibit hyperproliferation at 4-weeks of age. However, proliferation returns to baseline by 6-weeks, which is earlier than females. Each dot represents an individual animal. Error bars represent mean ± s.e.m. Statistical analysis was performed using two-way ANOVA followed by Tukey’s multiple comparison test. Scale bars, 100µm.

Next, we assessed proliferation in *Znrf3* cKO adrenals. Previously, we observed hyperplasia at 6-weeks in females^18^. Consistent with these prior observations, the Ki67-index in female *Znrf3* cKOs was significantly higher than controls at both 4- and 6-weeks (Fig. 2c). However, proliferation was reduced to baseline control levels by 9-weeks. This striking drop in proliferation was coincident with the onset of adrenal regression we had observed by adrenal ultrasound and weight. Male *Znrf3* cKOs also displayed an initial phase of hyperplasia evident by the high rate of proliferation at 4-weeks. However, the Ki67-index returned to baseline by 6-weeks (Fig. 2d). These results indicate that adrenal cortex hyperplasia in *Znrf3* cKOs is not sustained. Further, the onset of regression occurs earlier in males as compared with females, suggesting an underlying sexual dimorphism.

### The phenotypic switch from hyperplasia-to-regression is induced by cell cycle arrest

To identify potential mechanisms that mediate the switch from hyperplasia to regression, we performed bulk RNA-seq on control and *Znrf3* cKO adrenals. We focused our analysis on females at 6-weeks (hyperplasia), 9-weeks (regression onset), and 12-weeks (regression). Control adrenals had a highly stable expression signature. Only 23 genes were differentially expressed between 6- and 9-weeks (17 up-regulated, 6 down-regulated, padj<0.05). In contrast, we observed a striking change in gene expression profile in *Znrf3* cKOs, with 720 significant differentially expressed genes (DEGs) (383 up-regulated, 337 down-regulated, padj<0.05) (Fig. 3a).

**Figure 3.**
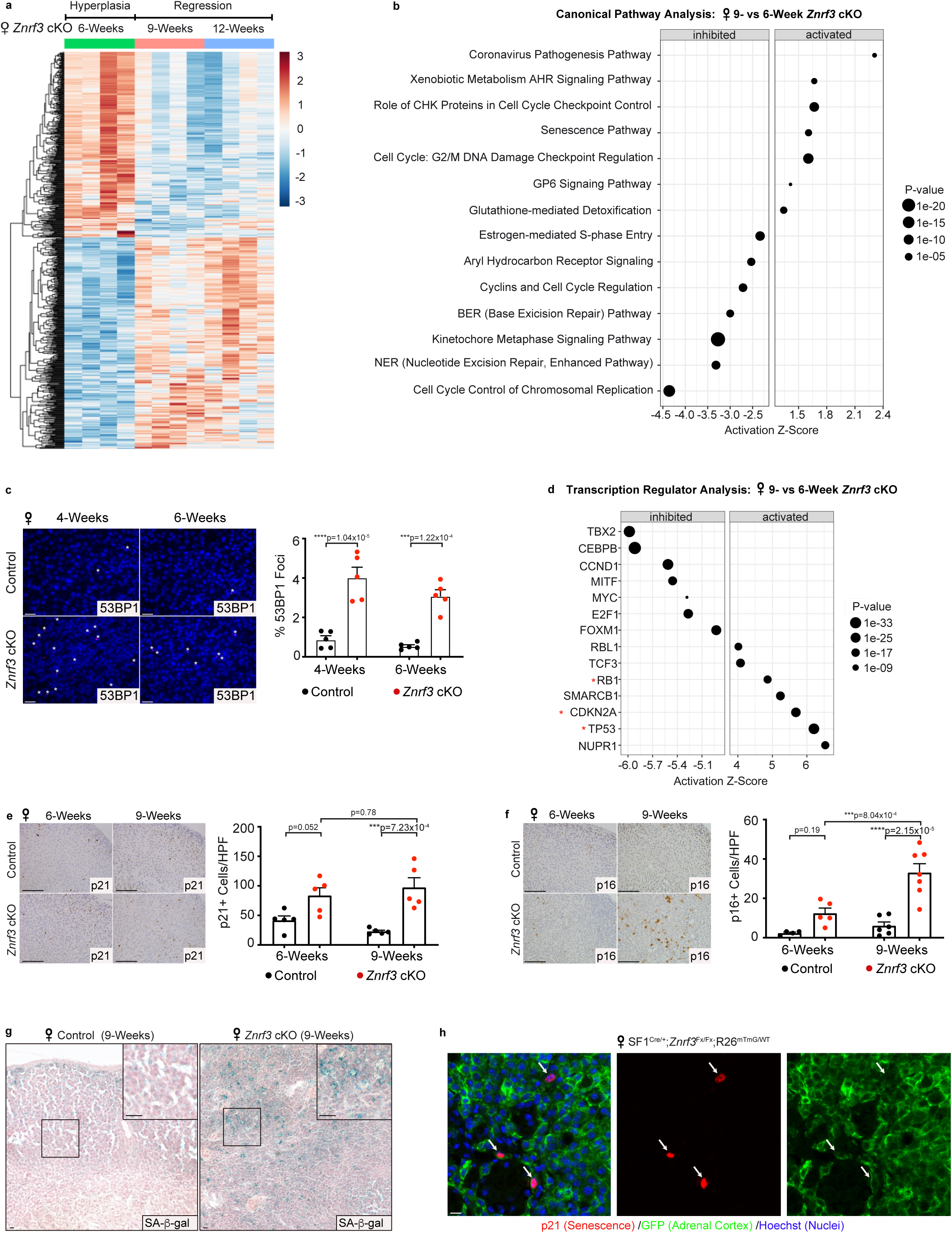
The switch from hyperplasia to regression is marked by activation of cellular senescence. (a) Bulk RNA-seq reveals a significant change in gene expression signature during the switch from hyperplasia to regression. Heatmap of significant DEGs in female *Znrf3* cKO adrenals at 6-, 9-, and 12-weeks of age (padj <0.05). (b) IPA identifies the most significantly altered canonical pathways in 9- vs 6-week female *Znrf3* cKO adrenals. (c) DNA damage as measured by 53BP1 foci is significantly increased in female *Znrf3* cKO adrenals compared to controls at 4- and 6-weeks of age. Quantification of 53BP1 foci was performed using QuPath digital image analysis based on the number of positive foci and normalized to total nuclei. Asterisks indicate 53BP1-positive foci. Scale bars, 10µm. (d) IPA of the most significantly activated or inhibited transcription regulators in 9- vs 6-week female *Znrf3* cKO adrenals. (e) p21, (f) p16^INK4a^, and (g) senescence associated beta-galactosidase (SA-β-gal) are significantly increased in 9-week female *Znrf3* cKO adrenals compared to controls. Scale bars, 100µm. Quantification of p21 and p16^INK4a^ IHC was performed using QuPath digital image analysis based on the number of positive cells per high powered field (HPF). Representative SA-β-gal images obtained from analysis of 3 independent mice are shown. (h) Co-staining of p21 and GFP in 9-week female *Znrf3* cKO mice containing the R26R^mT/mG^ lineage reporter tool confirms that p21-positive senescent cells are GFP-positive adrenal cortex cells. Representative images obtained from analysis of 5-female and 7-male animals are shown. Scale bars, 10µm. (b, d) IPA analysis includes 4 biological replicates per group. Statistical tests were performed using IPA, p<0.05. (c, e-f) Error bars represent mean ± s.e.m. Each dot represents an individual animal. Statistical analysis was performed using two-way ANOVA followed by Tukey’s multiple comparison test.

Using Ingenuity Pathway Analysis (IPA)^20^, we analyzed DEGs to identify canonical pathways associated with the phenotypic switch from hyperplasia to regression. Among the top inhibited pathways, we identified multiple pathways that regulate cell division, including cell cycle control of chromosomal replication and kinetochore metaphase signaling (Fig. 3b). Conversely, among the top activated pathways, we detected several signatures associated with enhanced DNA damage and cell cycle arrest. These included the role of CHK proteins in cell cycle checkpoint control and G2/M DNA damage checkpoint regulation (Fig. 3b). We validated these results *in situ* by measuring 53BP1, which is a DNA damage response factor that is recruited to DNA double-strand breaks^21,22^. Consistent with our RNA-seq data, we observed a significant increase in 53BP1 foci in *Znrf3* cKOs compared to controls (Fig. 3c, Extended Data Fig. 2a). Taken together, these data suggest that *Znrf3*-deficient adrenal cortex cells accumulate DNA damage and initiate cell cycle arrest near the onset of regression.

### Adrenal regression is associated with activation of cellular senescence

We next performed upstream regulator analysis (URA) to identify candidate factors that could coordinate the observed change in gene expression profile during the switch from hyperplasia to regression. URA is a novel IPA tool that analyzes the relationship between DEGs to identify potential upstream factors^20^. We focused our analysis on transcription regulators and identified 47 candidates (23 activated, 24 inhibited, p<0.05). Among the most significantly activated regulators, we found TP53 (Z-score=6.20), CDKN2A (Z-score=5.68), and RB1 (Z-score=4.86) (Fig. 3d), which are central regulators of cellular senescence – one of the top activated canonical pathways (Fig. 3b).

Cellular senescence is an aging-associated stress response that causes proliferative arrest of damaged cells^23^. Enhanced cellular stress, which can be induced by a wide-range of stimuli, activates the p53 and p16^INK4a^ tumor suppressor proteins leading to cell cycle inhibition through p21 and Rb. Based on our RNA-seq analysis in combination with observed phenotypic changes, we hypothesized that cellular senescence was activated in *Znrf3* cKOs during the transition from hyperplasia to regression. To test this hypothesis, we measured multiple hallmarks of senescence at 6- and 9-weeks. We observed increased expression of the cell cycle inhibitors p21 (Fig. 3e, Extended Data Fig. 2b) and p16^INK4a^ (Fig. 3f, Extended Data Fig. 2c), as well as accumulation of senescence-associated beta-galactosidase (SA-β-gal) (Fig. 3g, Extended Data Fig. 2d) in adrenals from 9-week *Znrf3* cKOs compared to controls. To confirm senescence activation in adrenal cortex cells, as opposed to the stromal or immune compartment, we performed co-staining for p21 and GFP in adrenal glands isolated from mice containing the R26R^mT/mG^ lineage reporter^24^. In these animals, the adrenal cortex is permanently labeled with GFP through SF1-Cre-mediated recombination. We found that all p21-positive cells were GFP-positive adrenal cortex cells at 9-weeks (Fig. 3h). Collectively, these data demonstrate that cellular senescence is activated in *Znrf3*-deficient adrenal cortex cells during the switch from hyperplasia to regression.

### Senescent adrenal cortex cells develop a senescence-associated secretory phenotype

One defining feature of cellular senescence that distinguishes it from quiescence and other forms of growth arrest is the senescence-associated secretory phenotype (SASP)^25^. The SASP comprises multiple families of secreted proteins including soluble signaling factors (*e.g.* growth factors, cytokines, chemokines), secreted proteases, and secreted insoluble proteins/ECM components. Although this phenotype is both temporally dynamic and heterogenous across tissue types/modes of senescence induction^26–28^, the SASP collectively acts as a potent pro-inflammatory stimulus to facilitate tissue remodeling.

To begin to define the SASP in the context of *Znrf3* loss in the adrenal cortex, we performed URA for activated cytokines and growth factors in our RNA-seq dataset. During senescence activation at 9-weeks, we identified IL1B (Z-score=2.96), IFNγ (Z-score=2.48), and TNFSF13 (Z-score=2.00) as the most highly induced cytokines (Fig. 4a, Extended Data Fig. 3a). By 12-weeks, additional members of the IL1 (IL1A, Z-score=2.38), IL6 (IL6, Z-score=2.15 and OSM, Z-score=3.15), and TNF family (TNF, Z-score=3.12 and TNFSF12, Z-score=2.37) were significantly activated (Fig. 4b, Extended Data Fig. 3b). Gene set enrichment analysis (GSEA) at both 9-weeks (Fig. 4c) and 12-weeks (Fig. 4d) further confirmed significant positive enrichment for genes associated with IL6/JAK/STAT3 signaling, chemokine signaling, the inflammatory response, and the innate immune system. Growth factor analysis additionally revealed significant induction of SASP-associated factors, including TGFB3^29^ and BDNF^30^ (Extended Data Fig. 3c-d).

**Figure 4.**
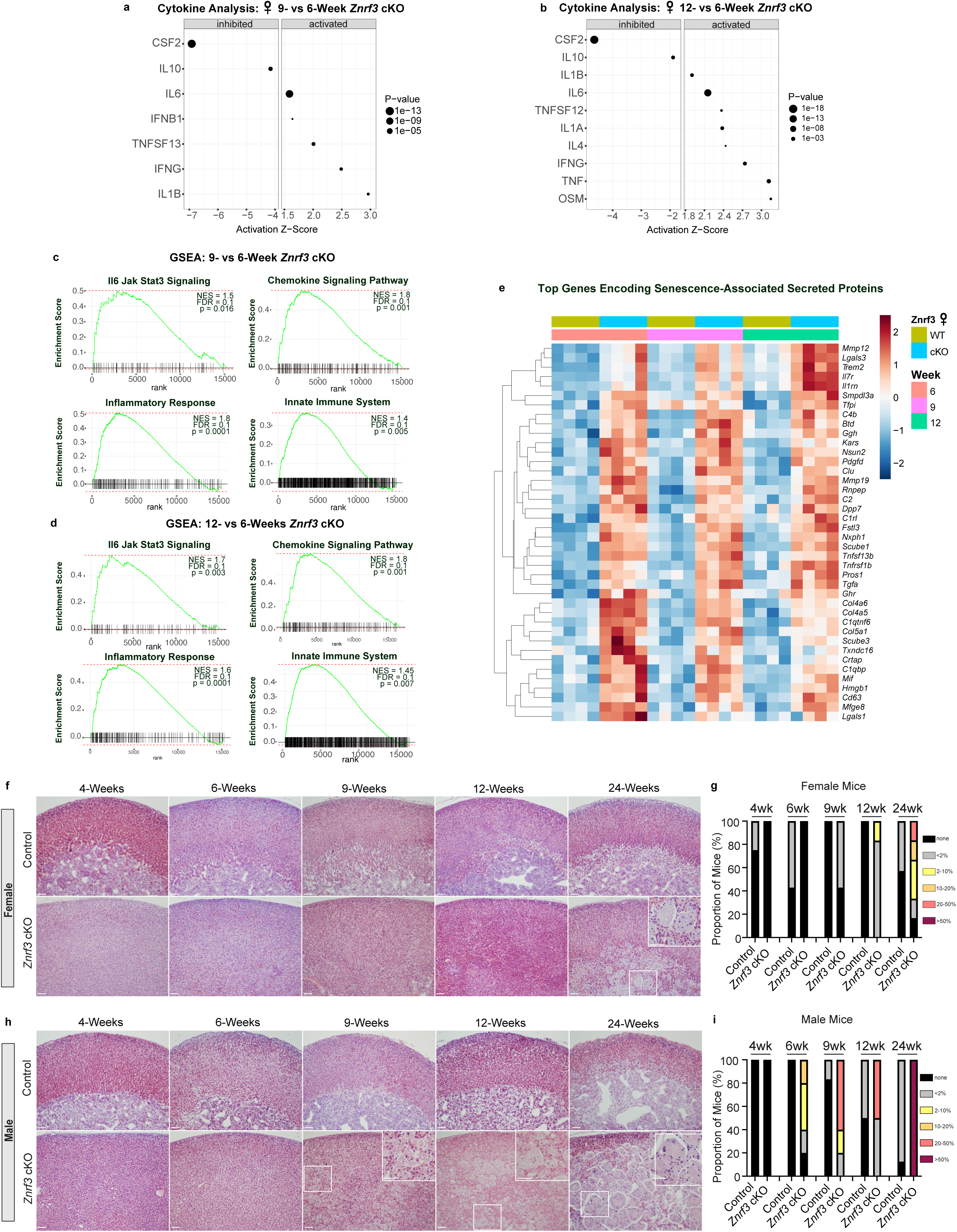
Senescent *Znrf3* cKO adrenal glands develop a functional SASP. (a) URA using bulk RNA-seq data predicts significantly activated and inhibited cytokines in 9-week and (b) 12-week *Znrf3* cKO adrenal tissue compared to 6-week. Data is representative of 4 biological replicates per group. Statistical tests were performed using IPA, p<0.05. (c) GSEA for inflammatory signatures (IL6/JAK/STAT3 signaling, chemokine signaling, the inflammatory response, and the innate immune system) identifies positively enriched pathways in 9-week and (d) 12-week *Znrf3* cKO adrenal tissue compared to 6-week. (e) Heatmap representing supervised hierarchical clustering of the 40 most significant DEGs that encode secreted proteins compared to controls. (f-g) Histological evaluation of female control and *Znrf3* cKO adrenal tissue based on H&E. Female *Znrf3* cKOs accumulate histiocytes (inset) between 12- and 24-weeks of age. (h-i) Histiocytes accumulate earlier and occupy a larger proportion of the adrenal gland in male *Znrf3* cKOs compared to females. Quantification was performed using QuPath digital analysis based on the proportion of histiocyte area normalized to total adrenal cortex area. Scale bars, 100µm.

Given significant activation of multiple SASP-associated cytokines and growth factors by URA, we next sought to more broadly characterize this phenotype and identify all genes that could potentially contribute to its composition. We analyzed expression of genes encoding secreted proteins^31^ that were significantly differentially expressed in *Znrf3* cKO adrenals compared to controls at 6-, 9-, and 12-weeks. We then prioritized top candidate SASP factors based on up-regulated genes that significantly changed during the phenotypic transition from hyperplasia to senescence activation/regression. This analysis revealed a set of early secreted factors with high induction at 6-weeks, and a set of later secreted factors whose expression progressively increased (Fig. 4e). We cross-referenced these early and late genes with the SASP Atlas^26^ and SeneQuest^23^, and found that >75% of factors we identified were established SASP components. These included early-response gene *Hmgb1*, which is a damage-associated molecular pattern molecule actively secreted upon cellular stress^26^ that initiates an IL6-dependent innate immune response^32,33^. Notably, *Hmbg1* is known to be secreted early after senescence induction, just preceding full SASP development^34^. Among our top late-response genes, we identified several central SASP components, including *Mmp12*^26,35–37^, *Lgals3*^38,39^, and *Il1rn*^37,40^, which play key roles in ECM remodeling and immune activation.

### SASP activity leads to significant immune cell recruitment in a sex-dimorphic manner

The SASP primarily functions to induce local inflammation and attract immune cells to clear out damaged, senescent cells^41,42^. Given significant SASP activity in *Znrf3* cKOs, we hypothesized immune cells would be recruited to the adrenal. We first tested this hypothesis by assessing broad histological changes using hematoxylin and eosin (H&E) staining. We performed H&E at each phenotypic stage from hyperplasia (4- and 6-weeks) to senescence activation (9-weeks) and regression (12- and 24-weeks). Strikingly, we observed accumulation of large multi-nucleated cells in *Znrf3* cKOs over time. These highly conspicuous cells first amassed in the innermost gland and became increasingly prevalent between 12- and 24-weeks in females (Fig. 4f-g). Upon pathologic review, we identified these as potential histiocytes, which are monocyte-derived immune cells known for their phagocytic role during tissue repair^43,44^. Interestingly, we observed an earlier and overall higher accumulation of these cells in males compared with females (Fig. 4h-i), consistent with the earlier onset of senescence and regression. Taken together, these results support the development of a functional adrenal SASP after prolonged *Znrf3* deficiency.

### scRNA-seq reveals innate and adaptive immune activation following cellular senescence

Histiocytes are complex structures derived from antigen-presenting dendritic cells (DCs) or phagocytic macrophages^43,44^. Moreover, histiocytes function in part to clear out neutrophils^45^. As a result, accumulation of histiocytes in *Znrf3* cKOs suggested that SASP-mediated tissue remodeling was likely driven by a multifaceted immune response. We performed scRNA-seq to provide an unbiased profile of the immune microenvironment following senescence. We specifically focused on the initial switch from hyperplasia (6-weeks) to regression (9-weeks) where we expected to identify the most proximal senescence-driven immune response. In total, we sequenced 18,294 cells from dissociated whole adrenal glands obtained from female *Znrf3* cKOs at 6- or 9-weeks. We then used unsupervised clustering methods to define groups of cells with similar transcriptional profiles and determined the identity of each cluster based on established cell-type specific markers in conjunction with Cluster Identity PRedictor (CIPR)^46^.

scRNA-seq revealed a total of 19 distinct cell clusters (Fig. 5a-b). Expression of cell type-specific markers for each cluster are provided in Extended Data Fig. 4-6. Within the major adrenal clusters, we observed a massive decrease in zona fasciculata cells between 6- and 9-weeks (Fig. 5c-d, Supplemental Table 1), consistent with adrenal cortex regression. Among the immune compartment, we observed increases in both myeloid and lymphoid lineages. The most striking myeloid changes included increased macrophages, conventional type 1 DCs (cDC1), and suppressive neutrophils. Additionally, we observed a distinct cluster of neutrophils uniquely present at 9-weeks. These cells expressed high levels of transcripts characteristic of immature neutrophils^47,48^, including *Chil3*, *Ltf*, *Mmp8/9*, and *Camp* (Extended Data Fig. 5). Within the lymphoid lineage, natural killer cells and B cells were the main increased populations. Overall, our scRNA-seq uncovered a complex immune response following senescence activation with a prominent role for myeloid cells.

**Figure 5.**
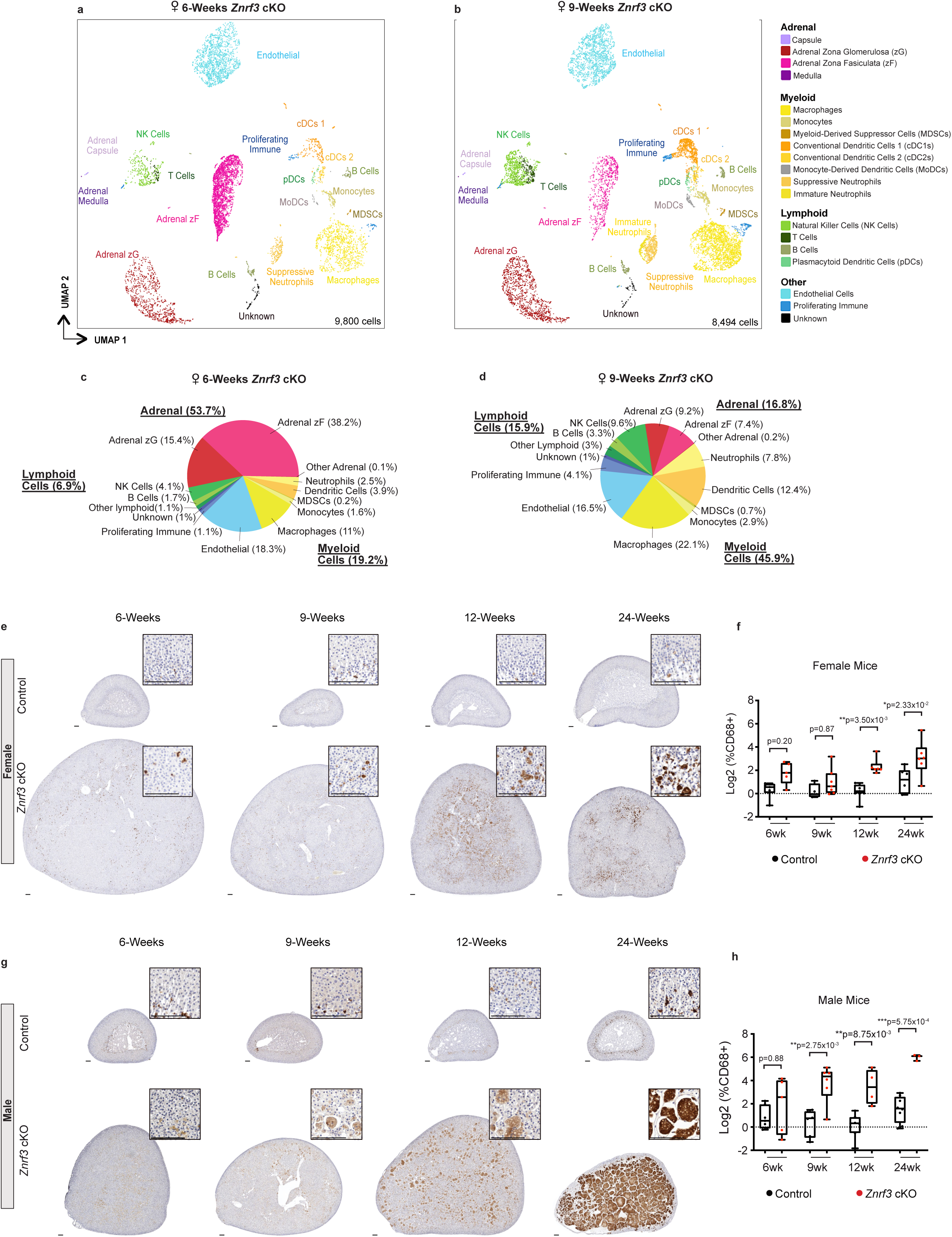
scRNA-seq reveals activation of the innate and adaptive immune system in response to cellular senescence and the SASP. UMAP plots with cluster identification of cell types in (a) 6-week (9,800 cells) compared to (b) 9-week (8,494 cells) female *Znrf3* cKO adrenal glands. Proportion of major (c-d) adrenal, myeloid, lymphoid, and other populations in 6-week and 9-week *Znrf3* cKO samples are graphed according to percentage of total cells. *In situ* validation of myeloid cell accumulation based on IHC for CD68 in control and *Znrf3* cKO adrenal tissue from (e-f) female and (g-h) male cohorts. Scale bars, 100µm. Quantification was performed using QuPath digital analysis based on the number of positive cells normalized to total nuclei. Each dot represents an individual animal. Box and whisker plots represent mean with variance across quartiles. Statistical analysis was performed on log2 transformed data using two-way ANOVA followed by Tukey’s multiple comparison test.

Given the large proportion of immune cells detected by scRNA-seq, we next performed IHC to measure *in situ* changes within the tissue microenvironment. We chose the broad myeloid marker, CD68, which was highly expressed across our DC, monocyte, neutrophil, and macrophage cell clusters (Extended Data Fig. 5). While we observed some heterogeneity between animals, CD68-positive myeloid cells progressively increased over time in female *Znrf3* cKOs compared to controls (Fig. 5e-f). In males, we observed a significant increase in CD68-positive cells at earlier stages compared with females. Moreover, CD68-positive cells occupied nearly the entire male gland by 24-weeks (Fig. 5g-h). These results suggest that cellular senescence triggers a robust immune response, which is more advanced in males, to mediate clearance of damaged adrenal cortex cells.

### Males exhibit a higher magnitude SASP

While both male and female *Znrf3* cKOs developed an inflammatory response following senescence activation, immune cells amassed earlier and to a greater extent in males. To determine whether this differential response was due to the SASP composition and/or magnitude, we performed bulk RNA-seq on male adrenals at 4- (hyperplasia), 6- (regression onset), and 9-weeks (regression). During the phenotypic switch from hyperplasia to regression (*i.e.* 4- to 6-weeks), we found 3,161 significant DEGs (1,558 up-regulated, 1,603 down-regulated, padj<0.05) (Fig. 6a). Mirroring what we observed in females, IPA identified several canonical pathways that regulate cell division as the most significantly inhibited (Fig. 6b). Moreover, cellular senescence was among the top activated pathways (Fig. 6b), and senescent mediators CDKN2A and TP53 were two of the top activated transcription regulators (Fig. 6c). Consistent with the strong immune phenotype and massive accumulation of phagocytic histiocytes in males, multiple immune-related signatures as well as phagosome formation were among the top activated pathways (Fig. 6b). This was similar to the later-stage response we observed in females at 12-weeks (Extended Data Fig. 7a). When we assessed the male cytokine profile (Fig. 6d, Extended Data Fig. 7b), we found substantial overlap with females, with 8 of the top 10 activated cytokines shared in both sexes. However, although the profile was similar, males consistently displayed a higher magnitude of induction, including IL1B (Z-score=5.92), IFNγ, (Z-score=7.54), and TNF (Z-score=6.83). This was further confirmed with enhanced GSEA enrichment of immune-related signatures (Fig. 6e, Extended Data Fig. 7c). These results suggest that both male and female *Znrf3* cKOs develop a SASP following senescence activation. However, males undergo the phenotypic switch earlier and display a higher magnitude of cytokine induction, ultimately leading to an enhanced immune response.

**Figure 6.**
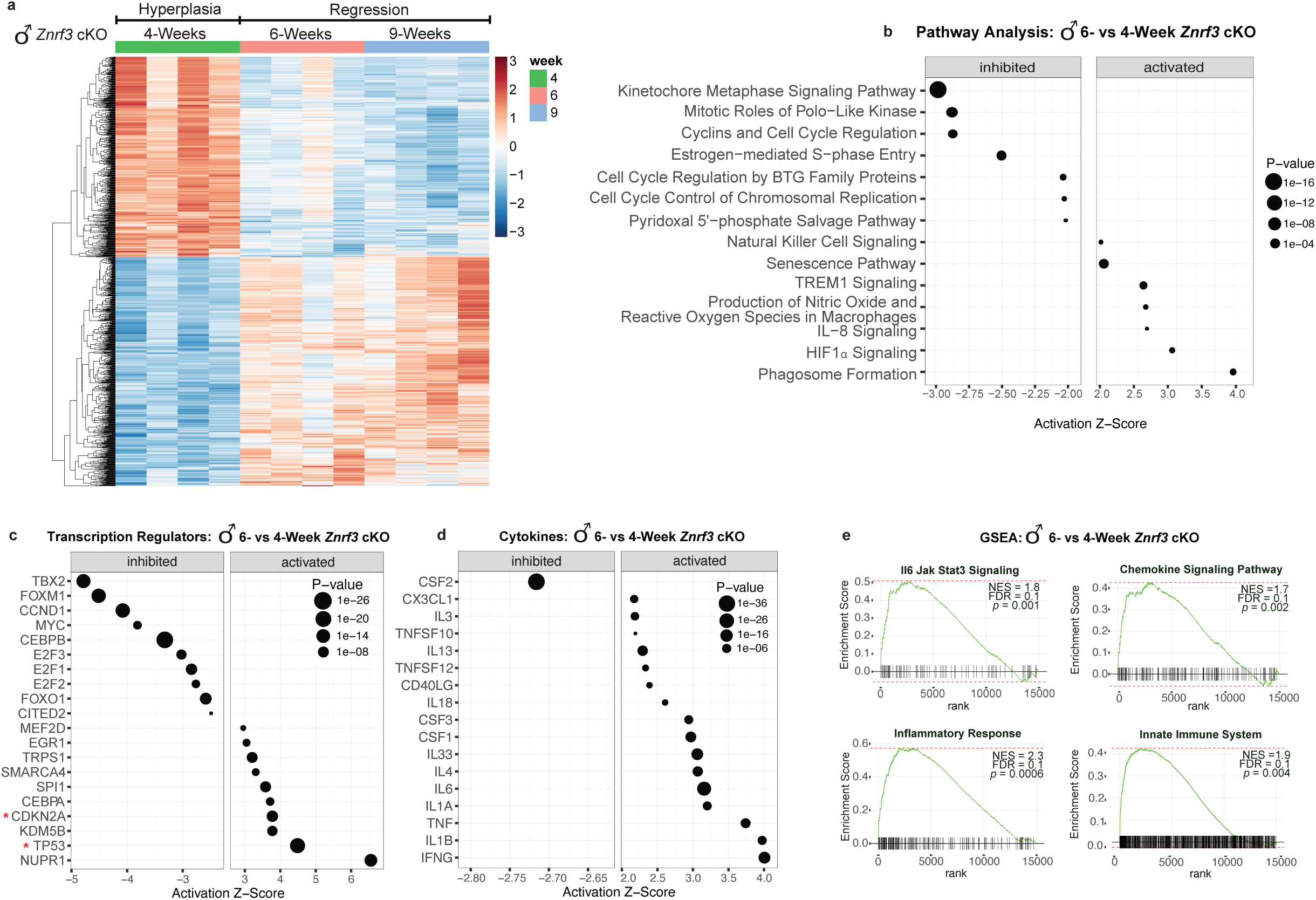
Senescence-mediated immune activation in male *Znrf3* cKO mice is characterized by higher SASP induction. (a) Heatmap of top 1,000 DEGs in *Znrf3* cKO adrenals at 4-, 6-, and 9-weeks of age (padj<0.05). IPA identified the top inhibited and activated (b) canonical pathways, (c) transcription regulators, (d) and cytokines in 6- vs 4-week male *Znrf3* cKO adrenals. Data is representative of 4 biological replicates per group and statistical tests were performed using IPA, p<0.05. (e) GSEA for inflammatory signatures (IL6/JAK/STAT3 signaling, chemokine signaling, the inflammatory response, and the innate immune system) identifies positively enriched pathways in 6- vs 4-week male *Znrf3* cKO adrenals.

### Following senescence-mediated tissue remodeling, a significant portion of mice develop adrenal tumors

Based on high levels of immune infiltration and adrenal degeneration, we predicted that *Znrf3* cKO animals would succumb to adrenal insufficiency. To test this hypothesis, we generated late-stage cohorts at 44-, 52- (1-year), and 78-weeks (18-months) (Fig. 7a-b). Surprisingly, rather than adrenal failure, a large proportion of *Znrf3* cKOs ultimately developed adrenal tumors (Fig. 7c). Notably, tumor development and progression were highly sex-dimorphic, with females exhibiting significantly increased malignancy compared with males.

**Figure 7.**
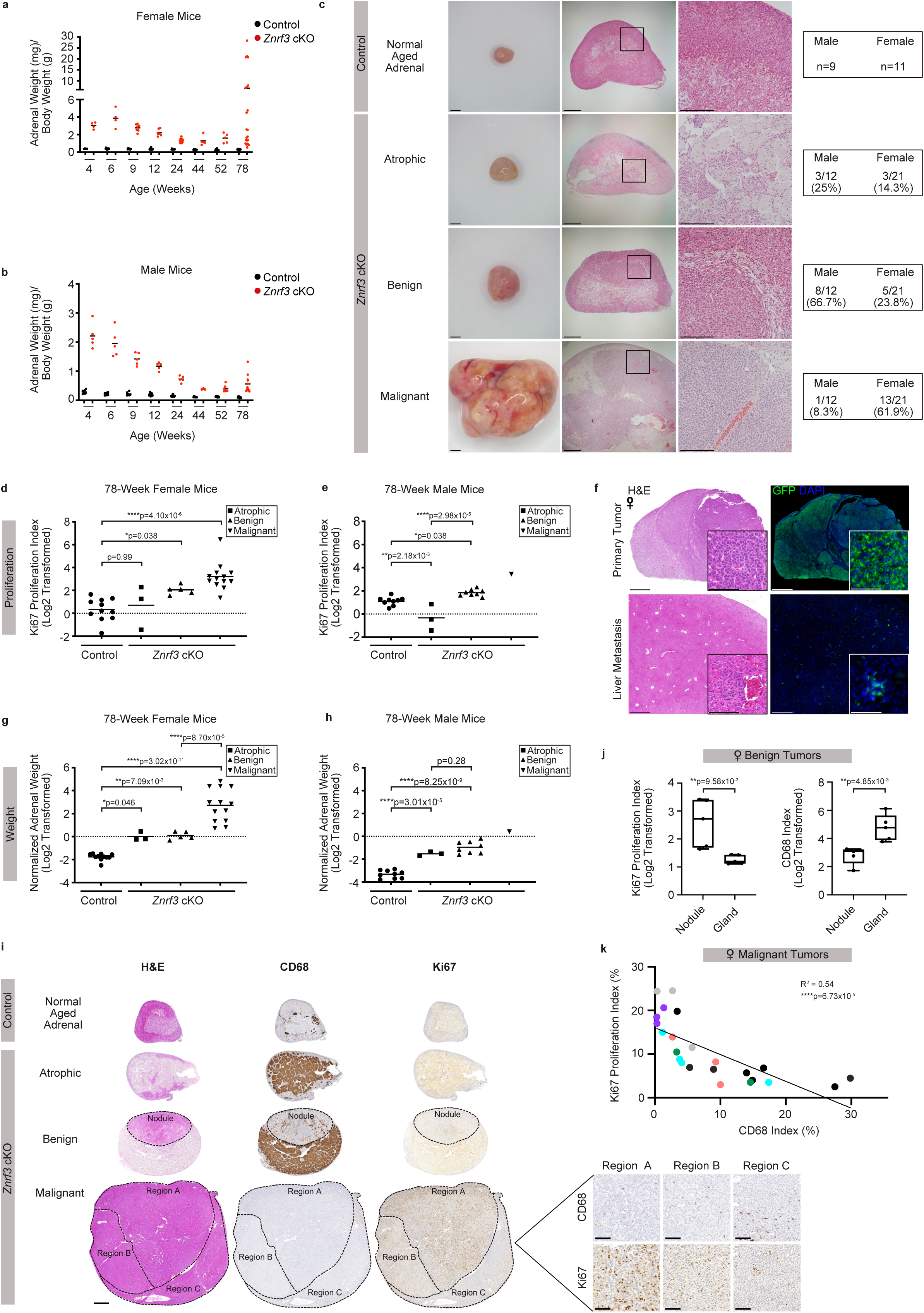
Following senescence-mediated remodeling of the tissue microenvironment, metastatic ACC tumors arise in a sex-dimorphic manner. Adrenal weight measurements in cohorts of (a) female and (b) male control and *Znrf3* cKO mice, including 44-, 52-, and 78-weeks of age. After 52-weeks, adrenal tumors of varying sizes begin to form. Each dot represents an individual animal. Black line indicates the mean. (c) Representative images of the gross histology (left) and H&E (middle & right) from each pathology observed at 78-weeks of age. Blinded samples were scored as atrophic, benign, or malignant. The proportion of each pathology in male compared with female cohorts is indicated. Scale bars, 1mm (left), 200µm (middle), 500µm (right). (d-e) Proliferation as measured by Ki67-index is significantly higher with progression to benign and malignant tumors in *Znrf3* cKO mice. Each symbol represents an individual animal. Black line indicates the mean. Statistical analysis was performed using one-way ANOVA followed by Tukey’s *post hoc*. (f) Malignant tumors metastasize to distant sites, including the liver. Representative images are shown from mice expression the R26R^mTmG^ lineage reporter, which permanently labels adrenal cortex cells with GFP. Scale bars, 100µm. (g-h) Tissue weight is significantly higher with progression to benign and malignant tumors in *Znrf3* cKO mice. Each symbol represents an individual animal. Black line indicates the mean. Statistical analysis was performed using one-way ANOVA followed by Tukey’s *post hoc*. (i) Representative H&E (left), CD68 (middle), and Ki67 (right) staining in atrophic, benign, and malignant *Znrf3* cKO adrenals from 78-week-old mice. Scale bars, 500µm (whole tissue), 100µm (high-magnification inset). (j) In *Znrf3* cKO adrenals, benign nodules have a significantly higher Ki67-index and lower CD68-index compared to the background gland. Each dot represents an individual nodule. Box and whisker plots represent mean with variance across quartiles. Statistical analysis was performed using two-tailed Student’s t-test. (k) In malignant tumors, there is a significant inverse correlation between the Ki67- and CD68-index. Each dot represents a distinct tumor region. Colors represent regions from the same tumor. Statistical analysis was performed using simple linear regression (n=23 tumor regions from 6 animals).

To unbiasedly evaluate tumors, we performed H&E and Ki67 staining. Blinded samples were then pathologically reviewed based on criteria derived from the Weiss system conventionally used to evaluate adrenal cortical neoplasms in adults^49,50^. This analysis revealed three main phenotypes in *Znrf3* cKO adrenals: atrophic glands, benign lesions, and malignant tumors (Fig. 7c). Atrophic glands were characterized by high histiocyte accumulation (Fig. 7c) and an overall Ki67-index at or below age-matched controls (Fig. 7d-e). This pathology was observed in 14.3% of females (3/21 mice) and 25% of males (3/12 mice). Benign tumors were most prevalent in males (66.7%, 8/12 mice). These lesions were characterized by formation of discrete nodules contained within the encapsulated adrenal cortex (Fig. 7c) with a significantly higher Ki67-index compared to matched controls (Fig. 7d-e). Finally, malignant tumors were most prevalent in females (61.9%, 13/21 mice). We conservatively classified tumors as malignant strictly based on the presence of metastases, which occurred to clinically relevant sites including the lung and liver (Fig. 7f). However, malignant tumors also had a significantly higher Ki67-index (>10%) (Fig. 7d-e), were significantly larger in size (Fig. 7g-h), and contained Weiss features of ACC (data not shown). Overall, these findings demonstrate that *Znrf3* loss is permissive for ACC with advanced aging, in a sex-dependent manner.

### Myeloid cells are depleted with advanced tumor progression and predict patient outcome

Our *Znrf3* cKO model presented two striking sexual dimorphisms. Males exhibited an earlier, more robust immune response following senescence activation and were less likely to develop malignant tumors with age. Conversely, females displayed a dampened senescence-induced immune response and were significantly more likely to develop large, metastatic tumors. To further examine the relationship between the immune microenvironment and tumor progression, we quantified CD68-positive myeloid cells in atrophic, benign, and malignant tumors. Atrophic glands exhibited a high CD68-index, consistent with the accumulation of histiocytes noted on H&E (Fig. 7i, Extended Data Fig. 8a). In benign cases, there was an inverse correlation between proliferation and myeloid cell infiltration. Nodules displayed a significantly higher Ki67-index and lower CD68-index compared to the background gland (Fig. 7i-j, Extended Data Fig. 8b). This apparent exclusion of myeloid cells in more proliferative regions was also evident in malignant tumors. Here, we took advantage of high inter- and intra-tumoral heterogeneity and analyzed CD68-positive myeloid cells across different tumors as well as subpopulations. We found regions with the highest Ki67-index had the lowest CD68-index (R^2^=0.54, ****p=6.73×10^-5^) (Fig. 7i and 7k). Taken together, these results suggest that as adrenal tumors become more aggressive, the immune microenvironment is remodeled to exclude tumor-suppressive myeloid cells.

Given these observations, we next wanted to assess myeloid cell infiltration in ACC patient tumors. A myeloid response score (MRS) was recently established in hepatocellular carcinoma (HCC) that predicts patient prognosis and therapy response^51^. We evaluated expression of each HCC-MRS gene in our scRNA-seq dataset and identified 4 genes (*Cd33*, *Cd68*, *Itgam*, *Msr1*) that were myeloid-specific (Fig. 8a-d) and whose combined expression represented the full adrenal myeloid compartment (Fig. 8e). When then evaluated the adrenal MRS (AMRS) in TCGA-ACC tumors in conjunction with established markers of poor prognosis (High-*MKI67*^52^ and Low-*G0S2*^53^) (Fig. 8f). Consistent with our *Znrf3* cKO model, females had a significantly lower AMRS compared with males (Fig. 8g). Moreover, Low-AMRS was associated with significantly reduced overall (Fig. 8h) and progression-free survival (Fig. 8i), even in females alone (Extended Data Fig. 9a-b). These results reveal an important role for myeloid cells in restraining ACC tumor progression with significant prognostic value.

**Figure 8.**
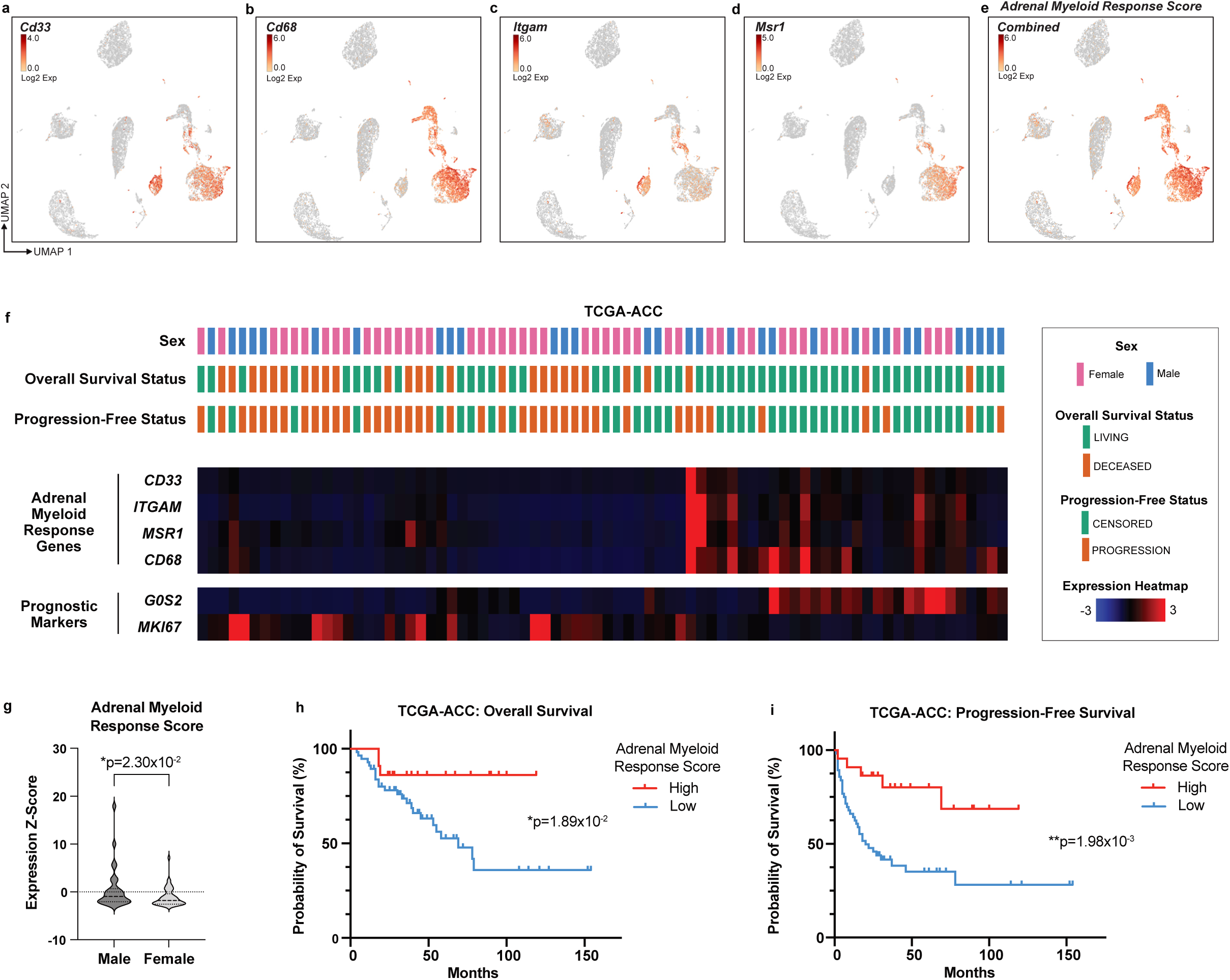
A low adrenal myeloid response score is associated with worse patient outcome in ACC. UMAP of scRNA-seq data from 6- and 9-week female *Znrf3* cKO mice. Expression of individual myeloid cell markers (a) *Cd33*, (b) *Cd68*, (c) *Itgam* (also known as CD11b), and (d) *Msr1* (also known as CD204) are shown. (e) Combined expression of these 4 markers was used to generate an adrenal myeloid response score (AMRS). (f) Analysis of TCGA-ACC data, including patient demographics and outcome, in accordance with mRNA expression of AMRS genes (*CD33, CD68, ITGAM, MSR1*) and established prognostic markers (High-*MKI67* (proliferation) and Low-*G0S2* (recurrent disease)). (g) AMRS is significantly lower in female compared to male TCGA-ACC patients, and low-AMRS is associated with shorter (h) overall and (i) progression-free survival. Statistical analysis was performed using one-tailed Mann-Whitney (g) or Log-rank Mantel-Cox test (h, i).

## Discussion

Our data demonstrate that loss of Wnt inhibitor *Znrf3* is permissive for metastatic adrenal cancer with advanced aging. However, genetic loss of *Znrf3* is not completely sufficient for tumor initiation and progression. The tumorigenic potential of *Znrf3* loss instead depends on extrinsic factors, including sex and the aged immune-microenvironment. These key factors are closely linked to cellular senescence, an age-associated stress response activated in *Znrf3*-deficient adrenals. Our *in vivo* characterization of senescence-induced tissue remodeling reveals a prominent role for myeloid-derived immune cells, including macrophages, DCs, and neutrophils, during the phenotypic switch from hyperplasia to regression. Interestingly, males activate cellular senescence earlier and mount a greater immune response, ultimately resulting in a lower incidence of metastatic tumors. We translate these findings to human ACC patients, where a high myeloid response score is associated with better outcome. Collectively, our model recapitulates the age- and sex-dependence of ACC and reveals a novel role for myeloid cells in restraining adrenal cancer progression with significant prognostic value.

Sex-specific differences in stem cell renewal, the tissue microenvironment, and the innate and adaptive immune response have been increasingly acknowledged as key factors in both aging^54^ and cancer^55^, including response to cancer immunotherapy^56^. Going forward, a key challenge in our model is to understand why males exhibit an advanced senescent response and subsequently become more protected from malignancy. While there are multiple underlying differences between males and females, a particularly appealing explanation is the role of male androgens, which have recently been shown to repress recruitment and proliferation of adrenal cortex stem cells^57^. These findings suggest that the renewal capacity of the male adrenal cortex is inherently lower at baseline compared with females, which could explain the earlier proliferative arrest in males. Additionally, androgens have independently been shown to activate cellular senescence *in vitro*^58,59^ and the androgen receptor (AR) can directly activate p21 through an androgen response element in the proximal promoter^60^. Consequently, androgen signaling in males may help accelerate cellular senescence through parallel pathway activation. Additionally, sex hormones, including both male androgens and female estrogen and progesterone, can elicit a range of effects on innate immune function. Most relevant to our studies, AR activity amplifies the NF-κb cistrome^61^, while estradiol and progesterone suppress LPS-induced cytokine production, including TNFA, IL1, and IL6^62,63^. Thus, the sex-dependent immune response in our model is likely determined, at least in part, by both pro-inflammatory effects of androgens combined with anti-inflammatory effects of estrogen and progesterone. This will be an important area of investigation moving forward.

Following extensive immune remodeling and advanced aging, *Znrf3* cKOs ultimately develop adrenal tumors at 12-18 months. This corresponds to peak ACC incidence at age 45-55^7,8^. Moreover, metastatic tumors in our model are significantly more common in females, which is analogous to the higher prevalence of ACC in women^7^. In *Znrf3* cKOs, tumor development requires escape from senescence-mediated tissue degeneration. While our studies focus on *Znrf3* as a common ACC tumor suppressor, we expect that tumorigenesis will be influenced by additional genetic mutations, particularly within genes that bypass cellular senescence that are recurrently altered in patients (*e.g. TP53*, *CDKN2A*, *RB1*, and others^10,11^). However, in addition to mutational events, we also expect that age-dependent changes in the tissue microenvironment significantly impact the functional consequence of acquired mutations. In particular, we observe a striking exclusion of myeloid cells with advanced tumor progression. These findings translate to human patients and align with the immune-poor nature of ACC^64^ as well as the modest efficacy of immunotherapy thus far^65^. However, given that a strong intra-tumor IFNγ signature correlates with better immunotherapy response in other cancers^66–68^, exploiting underlying mechanisms that regulate immune cell recruitment in our model may ultimately help enhance the efficacy of immune-based therapy for ACC and other immune-cold tumors.

Overall, our *Znrf3* cKO model is a highly unique *in vivo* system that highlights the pleiotropic effects of cellular senescence in cancer. Traditionally, cellular senescence has been studied as a tumor suppressive mechanism that halts proliferation of damaged cells at risk for neoplastic transformation^69^. However, more recent studies reveal that some senescent cells may be released from their growth arrest and actually promote more aggressive phenotypes long-term^70,71^. In our model, tumors may potentially arise from cells that *evaded* senescence or from cells *released* from senescence. Newly developed lineage tracing tools to track senescent cells *in vivo* will provide further insight on the origin of these tumors as well as metastatic outgrowths. This has important clinical implications given that conventional chemotherapy agents, including EDP (etoposide, doxorubicin, and cisplatin) frequently used as standard-of-care for advanced ACC^72^, are known to act in part by triggering ‘therapy-induced senescence’ (TIS)^73^. Consequently, EDP and other forms of TIS may favor long-term maintenance of senescent tumor cells primed for late-stage recurrence^74^. Senolytic agents that selectively kill senescent cells may be a promising strategy to eliminate tumor cells following TIS, which is an active area of ongoing clinical investigation^75^ and future study.

## Methods

### Mice

All animal procedures were approved by the Institutional Animal Care & Use Committee at the University of Michigan and University of Utah. Mouse strains used in this study have been previously described: SF1-Cre-high^17^, *Znrf3*-floxed^13^, R26R-mTmG^24^. All animals were maintained on the C57Bl/6J background with a 12-h light/12-h dark cycle and ad lib access to food and water. Littermate control animals were used in all experiments.

### Ultrasound

Mice were anesthetized using 3% isoflurane in O_2_ (1 liter per minute) in an induction chamber until reaching the surgical plane. Following induction, mice were weighed and moved to a heated platform in the prone position in a modified Trendelenburg. To maintain a surgical plane of anesthesia, 1–1.5% isoflurane in O_2_ was supplied via nose cone. Eye lubricant was applied to prevent corneal damage during prolonged anesthesia. ECG and respirations were monitored via non-invasive resting ECG electrodes. Hair was removed from the left dorsal area with depilatory cream. Two-dimensional (B-mode) and three-dimensional ultrasound images were recorded using a VisualSonics Vevo-2100 *in vivo* micro-imaging system with a MS 550D transducer, which has a center frequency of 40 MHz and a bandwidth of 22-55MHz.

### Adrenal weight

Adrenals were cleaned of excess fat, weighed, and snap frozen in liquid nitrogen or fixed as described for histology. In control animals, we observed an age-dependent increase in adrenal weight across our time series that significantly correlated with an age-dependent increase in body weight (Extended Data Fig. 10a-c). To account for these normal aging differences, adrenal weights in control and *Znrf3* cKO mice were compared based on the adrenal-to-body weight ratio calculated as the sum of both adrenal glands (mg) normalized to body weight (g). Raw adrenal weight data is provided for reference in Extended Data Fig. 10d-e.

### Histology

Adrenals were fixed in 10% normal buffered formalin (Fisher Scientific, 23-427098) for 24 hours at room temperature, paraffin-embedded, and cut into 5µm sections. Antibody information and staining conditions are listed in Supplementary Table 2. Endogenous peroxidase activity was blocked with 0.3% H_2_0_2_ for 30 minutes at room temperature. For brightfield IHC, primary antibodies were detected with HRP polymer solution (Vector Laboratories, MP-7401 (anti-rabbit), MP-7402 (anti-mouse), or MP-7405 (anti-goat)) and DAB EqV peroxidase substrate (Vector Laboratories, SK-4103-100), nuclei were counterstained with hematoxylin (Sigma, GHS132), and slides were mounted using Vectamount (Vector Laboratories, H-5000-60). For immunofluorescence (IF), primary antibodies were detected with HRP polymer solution and Alexa fluor tyramide reagent (Thermo Fisher, B40953 (488), B40955 (555), or B40958 (647)), nuclei were counterstained with Hoechst (Thermo Fisher, H3570), and slides were mounted using ProLong Gold (Thermo Fisher, P36930). Imaging was performed on a Zeiss Apotome microscope with an AxioCam MRm camera or a Pannoramic MIDI II (3DHISTECH) digital slide scanner. Quantification was performed using QuPath digital pathology software (Version 0.3.0).

### RNA isolation

Adrenals isolated from mice at the indicated time points were cleaned of excess fat, weighed, snap frozen in liquid nitrogen, and stored at -80°C. For homogenization, RLT lysis buffer (Qiagen, 74104) containing 1% β-mercaptoethanol was added to thawed adrenal tissue in high impact zirconium 1.5mm bead tubes (Benchmark Scientific, D1032-15). Samples were homogenized for 30 seconds using the Beadbug™ 3 (Benchmark Scientific, D1030), returned to ice, and homogenized for an additional 30 seconds. Total RNA isolation was completed using the RNeasy® Kit (Qiagen, 74104) according to manufacturer guidelines. DNases were removed using the RNase-Free DNase Set (Qiagen, 79254) and subsequently purified using the RNeasy MinElute Cleanup Kit (Qiagen, 74204) according to manufacturer guidelines.

### Bulk RNA-seq

Following RNA isolation and DNAse treatment, RNA quality was evaluated using an Agilent 2100 Bioanalyzer (Agilent Technologies). All samples used for sequencing had an RNA integrity number value >8.0. Libraries were prepared using the NEBNext Ultra II Directional RNA Library Prep with poly(A)mRNA Isolation protocol and sequenced using the NovaSeq Reagent Kit v1.5 150×150 bp sequencing protocol. Libraries were sequenced on an Illumina Novaseq 6000. The mouse GRCm38 genome and gene feature files were downloaded from Ensembl release 102 and a reference database was created using STAR version 2.7.6a with splice junctions optimized for 150 base pair reads^76^. Optical duplicates were removed from the paired end FASTQ files using clumpify v38.34 and reads were trimmed of adapters using cutadapt 1.16^77^. The trimmed reads were aligned to the reference database using STAR in two pass mode to output a BAM file sorted by coordinates. Mapped reads were assigned to annotated genes using featureCounts version 1.6.3. Expected gene counts were filtered to remove features with zero counts and features with fewer than 10 reads in any sample. The age and genotype were combined into a single column in the design formula and differentially expressed genes were identified using a 5% false discovery rate with DESeq2 version 1.30.1^78^. Male and female datasets were analyzed in separate DESeq2 models. Pathways were analyzed using the fast gene set enrichment package^79^ and Ingenuity Pathway Analysis (IPA)^20^. Z-scores reported from IPA represent the activation Z-score, which are used to infer the activation states of pathways and upstream regulators.

### SA-β-galactosidase staining

Freshly isolated adrenal tissue was cryopreserved in optimal-cutting-temperature (OCT) compound. Cryosections (10µm) were stained using the Senescence β-Galactosidase Staining Kit (Cell Signaling Technology, 9860) according to the manufacturer’s protocol with minor modifications. Briefly, cryosections were air-dried at room temperature for 30 minutes, fixed for 15 minutes, and washed with PBS. Sections were stained inside a sealed, humidified chamber at 37°C overnight. Sections were then counterstained with eosin, dehydrated, and mounted for imaging.

### Adrenal gland dissociation

Adrenal glands were obtained from 6- or 9-week-old mice following rapid decapitation in order to minimize stress-induced transcriptional changes. Adrenals were placed immediately into ice-cold 1X Hank’s Balanced Salt Solution (HBSS) containing calcium and magnesium (Thermo Fisher, 14025134). Tissues were then finely chopped and digested enzymatically using enzymes and reagents from the Neural Tissue Dissociation Kit (Mitenyi Biotec, 130-092-628). All steps were carried out at 4°C, including centrifugation. All tubes and pipette tips used to handle cell suspensions were pre-coated with 3% BSA in HBSS to prevent cell loss. During the dissociation, the cell suspension was gently agitated with mechanical pipetting every 10 minutes and visually assessed under a stereo microscope until the tissue was fully digested. The suspension was then filtered through 70µm filters to obtain a single cell suspension and enzymes were neutralized using HBSS containing 10% Fetal Bovine Serum (FBS). Red blood cells were removed using Red Blood Cell Lysis buffer (Roche, 11814389001) according to manufacturer guidelines and the cells were washed twice in HBSS containing 2% FBS before counting on a Countess Automated Cell Counter (Thermo Fisher).

### scRNA-seq

Immediately following dissociation, cells were stained with 0.1 µg/mL DAPI and 5µM Vybrant DyeCycle Ruby (Invitrogen, V10309). Viable Ruby-positive, DAPI-negative single cells were sorted based on fluorescent properties using the BD FACSAria III Cell Sorter at the University of Utah Flow Cytometry Shared Resource. Cells were collected in 1XPBS (calcium- and magnesium-free) containing 0.04% BSA and kept on ice. Single-cell droplets with a target capture of 10,000 cells were immediately prepared on the 10X Chromium at the Huntsman Cancer Institute High-Throughput Genomics Shared Resource. Single-cell libraries were prepared using the Chromium Next GEM Single Cell 3’ Gene Expression Library Construction Kit v3.1 according to manufacturer instructions. Sequencing was performed on an Illumina NovaSeq 6000 using an S4 flow cell generating 150-base pair paired-end reads at a target depth of 300 million reads per sample. Raw data was converted to demultiplexed fastqs with 10X Cellranger mkfastq. Reads were mapped to the prebuilt mm10 reference distributed by 10X Genomics (version mm10-2020-A) and genes were counted with 10X Cellranger count. Samples were also aggregated into a single Loupe file with 10X Cellranger aggr. Filtering parameters for gene count range (>200 and <5,500), unique molecular identified (UMI) counts (>200 and <30,000), and mitochondrial transcript fraction (<15%) were used to remove low-quality cells.

### scRNA-seq analysis and cluster identification

For analysis of scRNA-seq clusters, major cell types were distinguished using graph-based and k-means methods using the Loupe Cell Browser Interface. For major cell clusters, core cell-type specific genes were used to validate initial clustering methods: Adrenal cortex (*Nr5a1*, also known as *Sf-1*), immune (*Ptprc*), endothelial (*Plvap*), and medulla *(Th*) (Extended Data Fig. 4). Validation of subclusters was completed by identifying defined transcriptional signatures from previously annotated mouse scRNA-seq datasets^80–82^ and established identification markers (Extended Data Fig. 4-6). Both inter- and intra-cluster comparisons were performed for cells of similar lineage. Final clustering was based on both gene expression patterns and the proportion of significant DEGs between respective clusters. Validation of immune-based clusters was completed using Cluster Identity PRedictor (CIPR) correlation analysis to compare novel clusters with the established ImmGen reference dataset^46^.

### Adrenal Myeloid Response Score

The relative adrenal myeloid response score (AMRS) in each tumor from TCGA-ACC was calculated using mRNA expression Z-scores obtained from cBioPortal (cBioPortal.org). A composite enrichment score was calculated based on combined expression of *CD33*, *CD68*, *ITGAM* (also known as CD11B), and *MSR1* (also known as CD204). TCGA-ACC tumors without mRNA expression data were excluded (n=14), resulting in final analysis of 31 male and 47 female tumors. A score cutoff >0 was used to classify AMRS-high vs AMRS-low tumors.

### Statistical Analysis

Statistical analysis was performed using R or GraphPad Prism 9. For comparison of two groups, a two-tailed Student’s t-test was performed. For more than two groups, one-way ANOVA followed by Tukey’s *post hoc* test was performed. When analyzing two independent variables, two-way ANOVA followed by Tukey’s *post hoc* was used. If normal distribution could not be assumed, a non-parametric analysis was used or data was log2 transformed prior to statistical analysis. A p-value < 0.05 was considered significant and exact p-values for each test are indicated.

## Acknowledgments

We thank Drs. Hans Clevers and Boo-Kyoung Koo for providing Znrf3-floxed mice, and the late Dr. Keith Parker for providing SF1-Cre transgenic mice. Research reported in this publication utilized the High-Throughput Genomics and Bioinformatic Analysis shared resource and the Biorepository and Molecular Pathology shared resource at Huntsman Cancer Institute at the University of Utah, and was supported by the National Cancer Institute of the National Institutes of Health under Award Number P30CA042014. We also utilized the University of Utah Flow Cytometry Shared Resource supported by the Office of the Director of the NIH Award Number S10OD026959 and NCI Award Number 5P30CA042014-24. The content is solely the responsibility of the authors and does not necessarily represent the official views of the NIH. We especially thank Drs. James Marvin and Opal Allen for respective technical assistance with flow cytometry and 10X Genomics library prep. We also thank Drs. Jay Gertz, Ben Myers, and Sheri Holmen for helpful scientific discussions and comments on the manuscript. This work was supported by funding from a Cancer Center Support Grant (P30CA040214, K.J.B), the V Foundation (V2021-021, K.J.B.), and 5 For The Fight (K.J.B.).

## Author Contributions

Conceptualization, K.M.W, G.D.H., K.J.B.; Experimentation, K.M.W., L.L., L.J.S., J.L.A., K.C., K.J.B; Bioinformatic analysis, K.M.W., B.K.L, C.J.S., H.A.E.; Pathologic review, T.J.G.; Data analysis, K.M.W, L.J.S., J.L.A., K.C., K.J.B.; Writing – Original Draft, K.M.W., K.J.B; Writing – Review & Editing, all authors.

---

**Supplemental Table 1.**
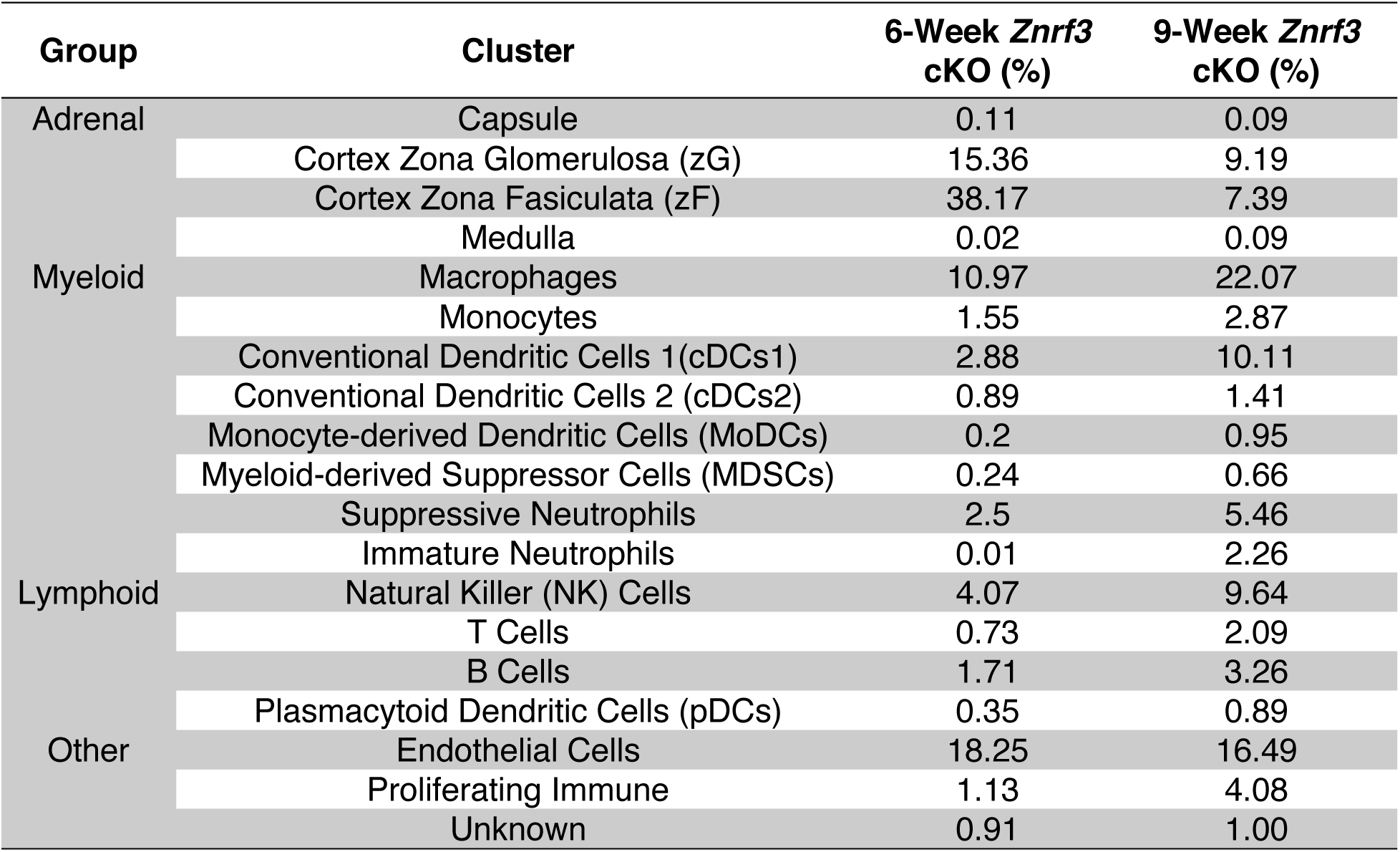
scRNAseq clusters in whole adrenals isolated from 6-week versus 9-week female *Znrf3* cKO mice.

**Supplemental Table 2.** List of antibodies and staining conditions for immunohistochemistry.

| **Target** | **Company, Catalog #** | **Species** | **Unmasking** | **Dilution** | **Blocking Buffer** | **Antibody Diluent** |
| --- | --- | --- | --- | --- | --- | --- |
| Cleaved caspase 3 | Cell Signaling, 9664 | Rabbit | 10mM NaCitrate, pH 6.0, 0.05% tween 20 | 1:200 | 2.5% HS + 1% BSA | 2.5% HS + 0.1% BSA |
| Ki67 | Thermo Fisher, MA5-14520 | Rabbit | 10mM NaCitrate, pH 6.0, 0.05% tween 20 | 1:200 | 2.5% HS | 2.5% HS |
| 53BP1 | Novus Biologicals, NB100-304 | Rabbit | 10mM NaCitrate, pH 6.0, 0.05% tween 20 | 1:1000 | 2.5% HS + 1% BSA | 2.5% HS + 0.1% BSA |
| p21 | Abcam, ab188224 | Rabbit | 10mM NaCitrate, pH 6.0, 0.05% tween 20 | 1:7500 | 2.5% HS + 1% BSA | 2.5% HS + 0.1% BSA |
| CDKN2A (p16<sup>INK4a</sup>) | Abcam, ab211542 | Rabbit | 10mM NaCitrate, pH 6.0, 0.05% tween 20 | 1:1000 | 2.5% HS + 1% BSA | 2.5% HS + 0.1% BSA |
| GFP | Abcam, ab5450 | Goat | 10mM NaCitrate, pH 6.0 | 1:1000 | 2.5% HS + 3% BSA | 2.5% HS + 1% BSA |
| CD68 | Abcam, ab125212 | Rabbit | 10mM NaCitrate, pH 6.0 | 1:1000 | 2.5% HS | 2.5% HS |

**Extended Data Figure 1.**
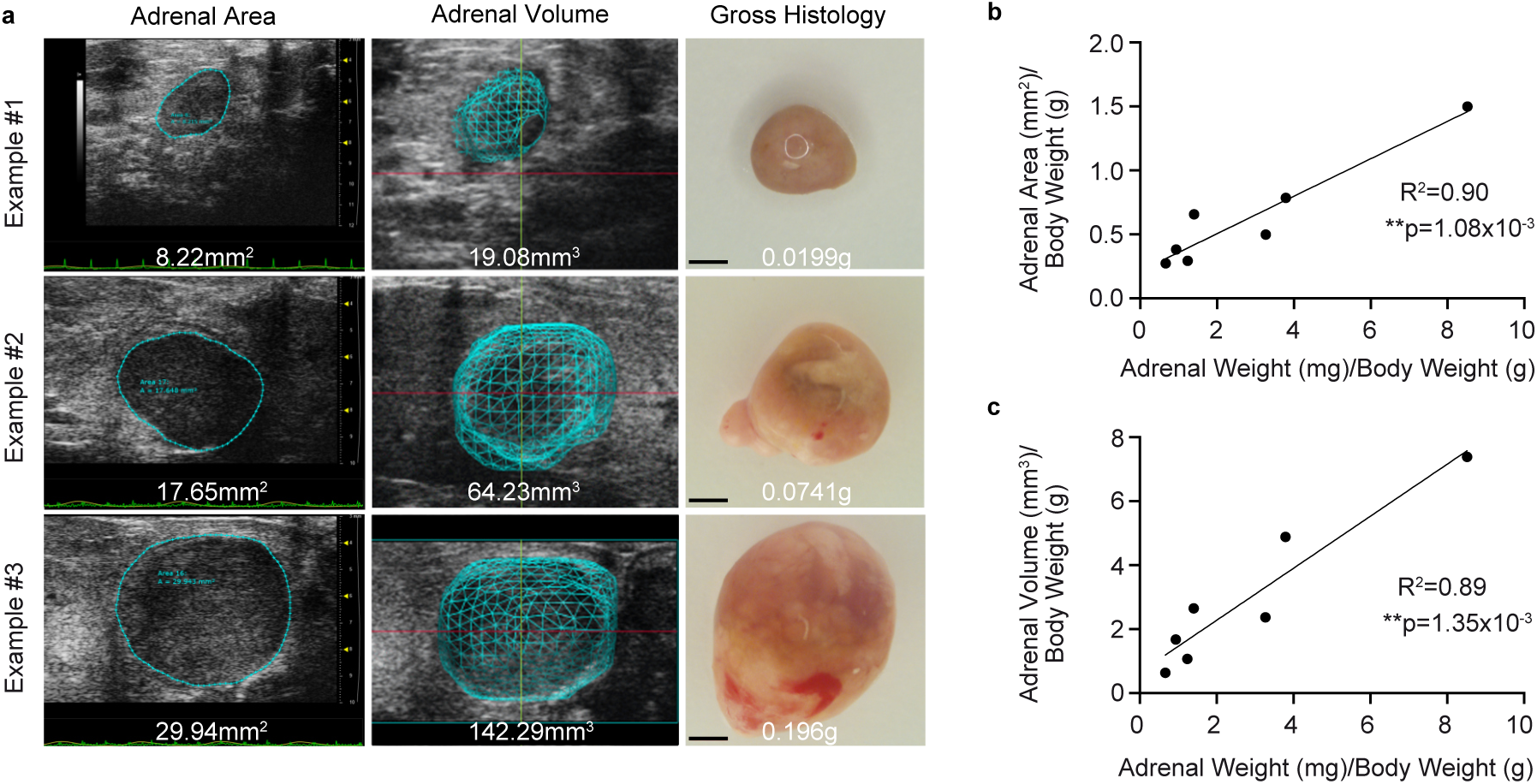
Ultrasound imaging provides an accurate measure of adrenal size in real-time. (a) Images of adrenal ultrasound area, ultrasound volume, and gross histology from representative animals with adrenals of varying size. Scale bars, 1mm. (b) Adrenal area and (c) adrenal volume are significantly correlated with adrenal weight. Ultrasound imaging was performed 24-hours prior to necropsy. All data shown is from female mice. Each dot represents an individual animal. Statistical analysis was performed using simple linear regression.

**Extended Data Figure 2.**
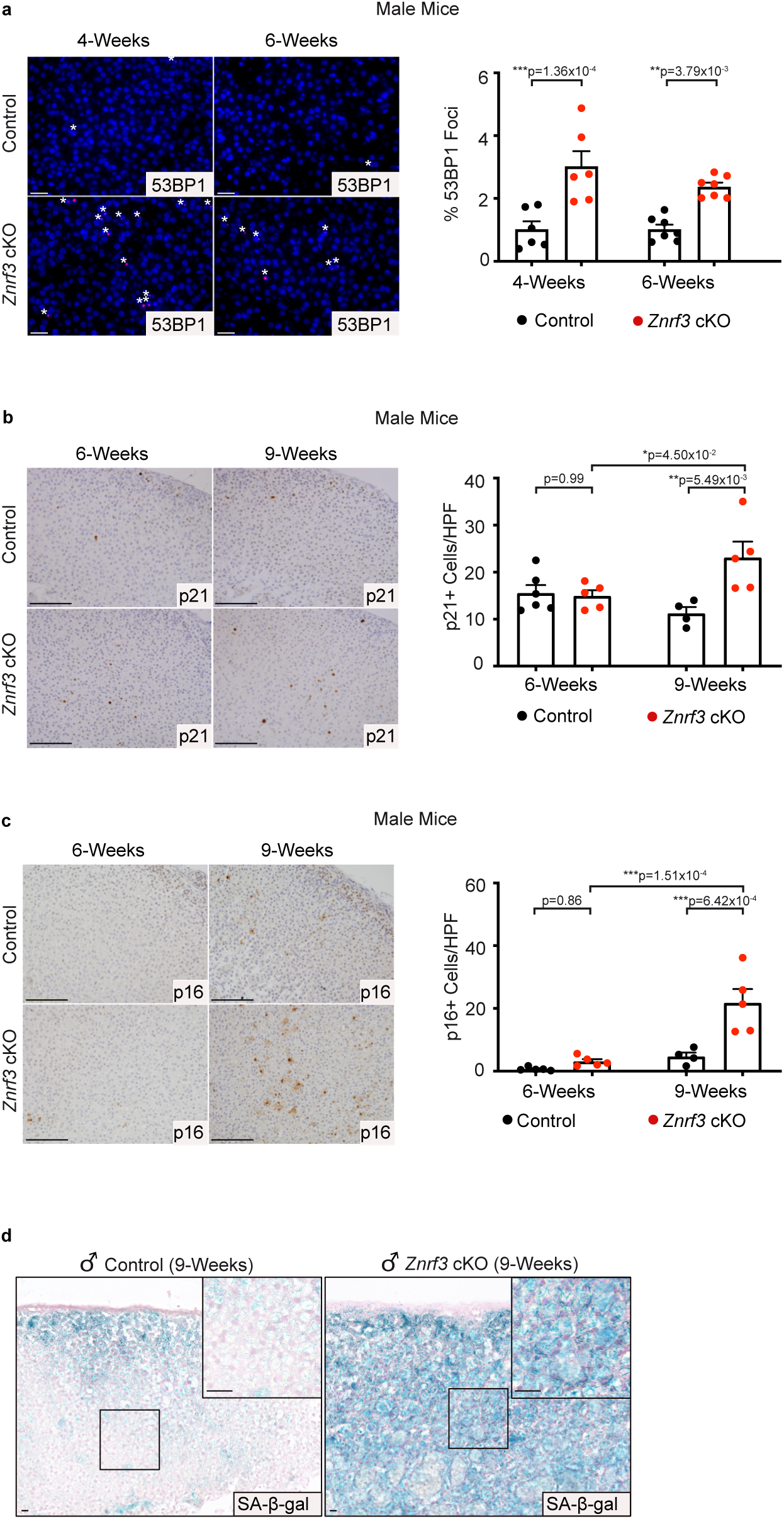
Male *Znrf3* cKO adrenals activate cellular senescence during adrenal regression. (a) DNA damage as measured by 53BP1 foci is significantly increased in male *Znrf3* cKO adrenals compared to controls at 4- and 6-weeks of age. Quantification of 53BP1 foci was performed using QuPath digital image analysis based on the number of positive foci and normalized to total nuclei. Asterisks indicate 53BP1-positive foci. Scale bars, 10µm. (b) p21, (c) p16^INK4a^, and (d) senescence-associated beta-galactosidase (SA-β-gal) are significantly increased in 9-week male *Znrf3* cKO adrenals compared to controls. Scale bars, 100µm. Quantification of p21 and p16^INK4a^ IHC was performed using QuPath digital image analysis based on the number of positive cells per high powered field (HPF). Representative SA-β-gal images obtained from analysis of 3 independent mice are shown. Error bars represent mean ± s.e.m. Each dot represents an individual animal. Statistical analysis was performed using two-way ANOVA followed by Tukey’s multiple comparison test.

**Extended Data Figure 3.**
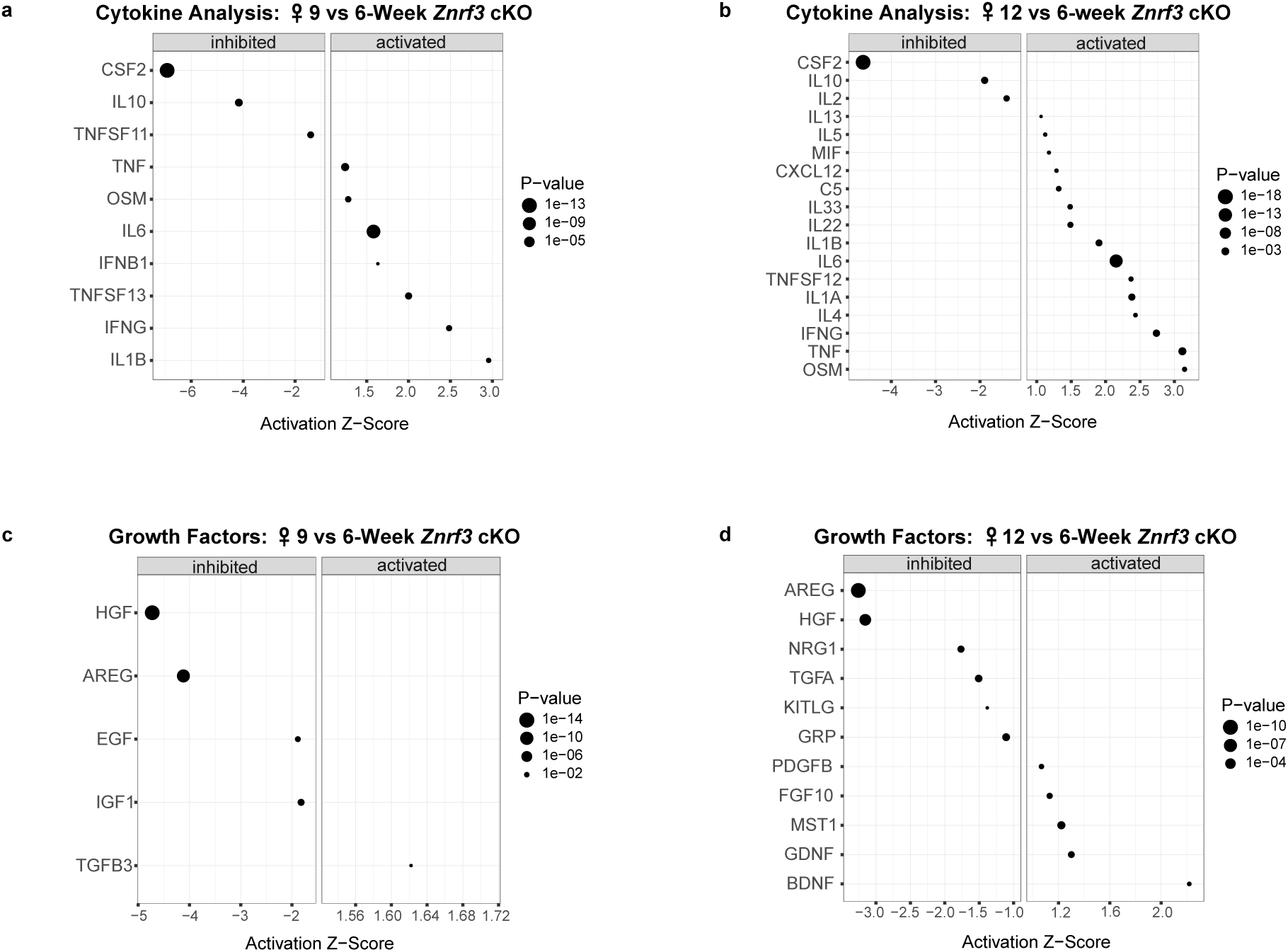
Senescent female *Znrf3* cKO adrenal glands activate production of cytokines and growth factors. (a) URA using RNA-seq data predicts significantly activated and inhibited cytokines in 9-week and (b) 12-week female *Znrf3* cKO adrenal tissue compared to 6-week. (c) URA predicts significantly activated and inhibited growth factors in 9-week and (d) 12-week female *Znrf3* cKO adrenal tissue compared to 6-week. Data is representative of 4 biological replicates per group. Statistical tests were performed using IPA, p<0.05.

**Extended Data Figure 4.**
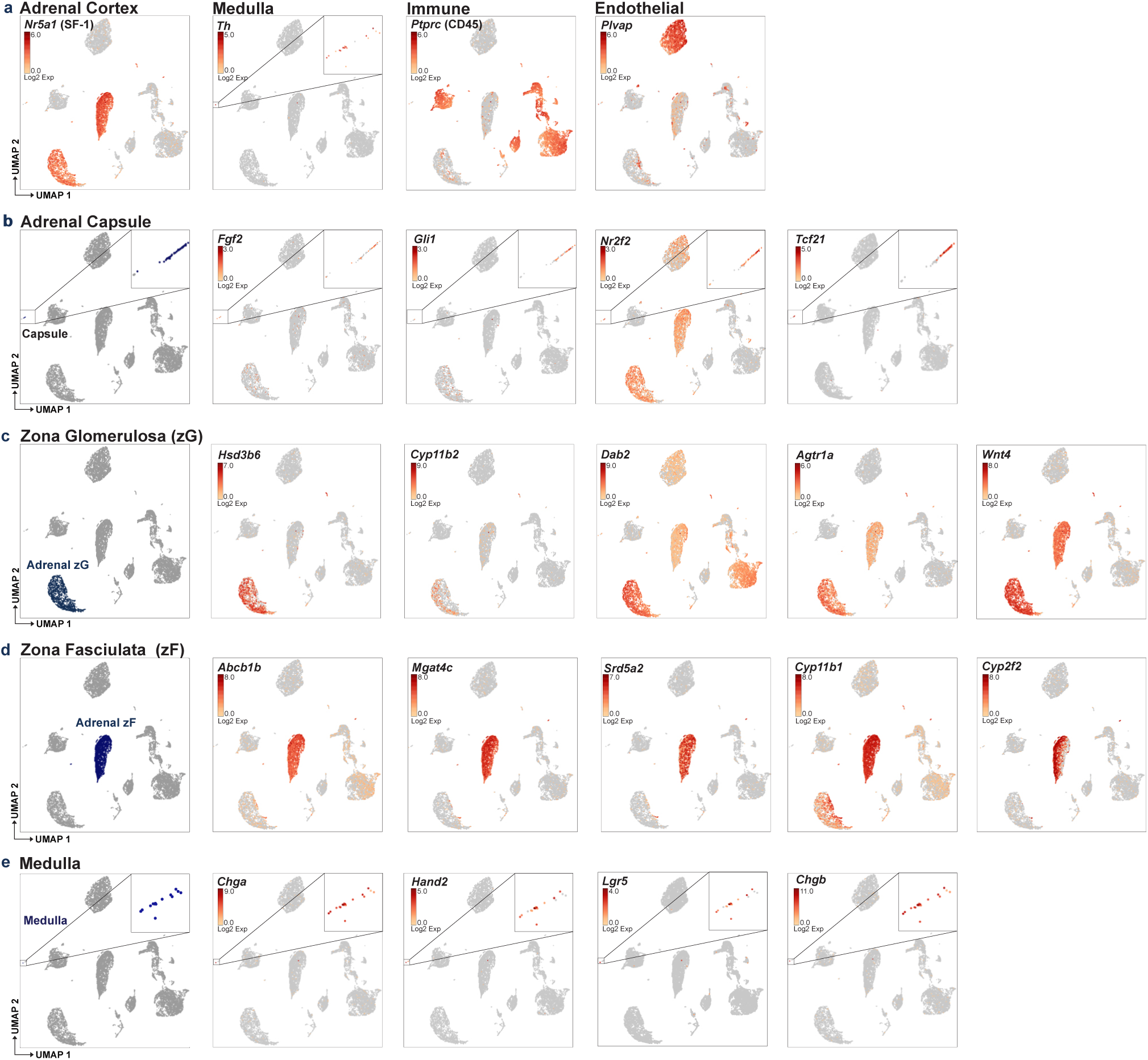
Validation of major cell types identified by scRNA-seq. (a) UMAP represents major clusters as identified by established cell-type specific marker genes: Adrenal cortex (*Nr5a1*, also known as SF-1), medulla (*Th*), immune (*Ptprc*, also known as CD45), and endothelial (*Plvap*). (b) Adrenal capsule identified by high expression of *Fgf2*, *Gli1*, *Nr2f2*, and *Tcf21*. (c) Zona Glomerulosa (zG) identified by high expression of *Hsd3b6*, *Cyp11b2*, *Dab2*, *Agtr1a*, *and Wnt4a*. (d) Zona Fasiculata (zF) identified by high expression of *Abcb1b*, *Mgat4c*, *Srd5a2*, *Cyp11b1*, and *Cyp2f2*. (e) Medulla identified by *Chga*, *Hand2*, *Lgr5*, and *Chgb*. Gene expression represented as log2 expression as indicated in respective UMAPs.

**Extended Data Figure 5.**
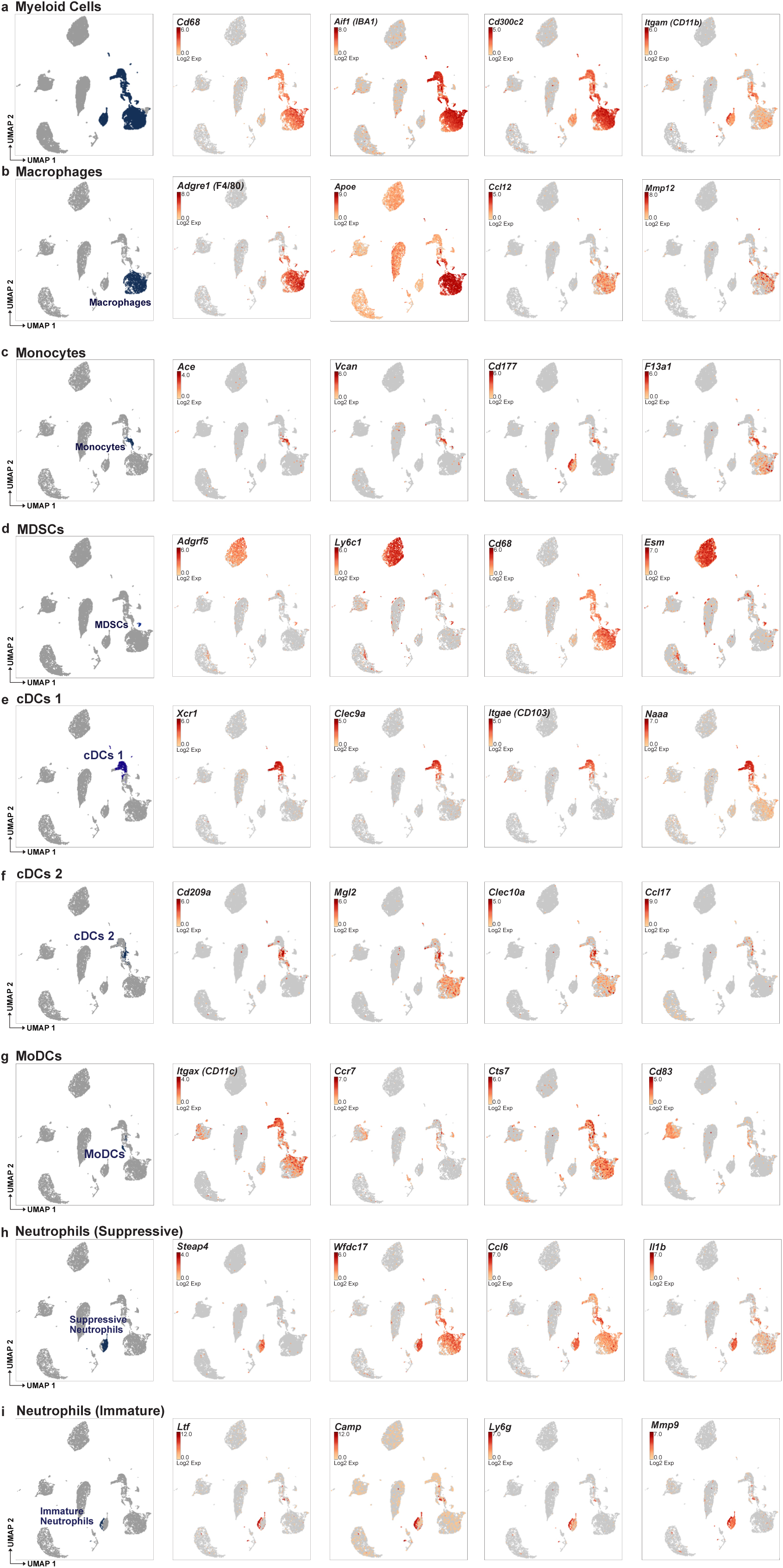
scRNAseq cluster validation of myeloid lineages. UMAP represents major clusters as identified by established cell-type specific marker genes. (a) Myeloid cells identified by high expression of *Cd68, Aif1* (also known as IBA1)*, Cd300c2,* and *Itgam* (also known as CD11b). (b) Macrophages identified by high expression of *Adgre1* (also known as F4/80)*, Apoe, Ccl12,* and *Mmp12*. (c) Monocytes identified by high expression of *Ace, Vcan, Cd117,* and *F13a1*. (d) MDSCs identified by high expression of *Adgrf5, Ly6c1, Cd68,* and *Esm.* (e) Conventional dendritic cells 1 (cDCs 1) identified by high expression of *Xcr1, Clec9a, Itgae* (also known as CD103), and *Naaa*. (f) Conventional dendritic cells 2 (cDCs 2) identified by high expression of *Cd2009a, Mgl2, Clec10a,* and *Ccl17*. (g) Monocyte-derived dendritic cells (MoDCs) identified by high expression of *Itgax* (also known as CD11c), *Ccr7*, *Cts7*, and *Cd83*. (h) Suppressive neutrophils identified by high expression of *Steap4*, *Wfdc17*, *Ccl6*, and *Il1b*. (i) Immature neutrophils identified by high expression of *Ltf*, *Camp*, *Ly6g*, and *Mmp9*. Gene expression is represented as log2 expression as indicated in respective UMAPs.

**Extended Data Figure 6.**
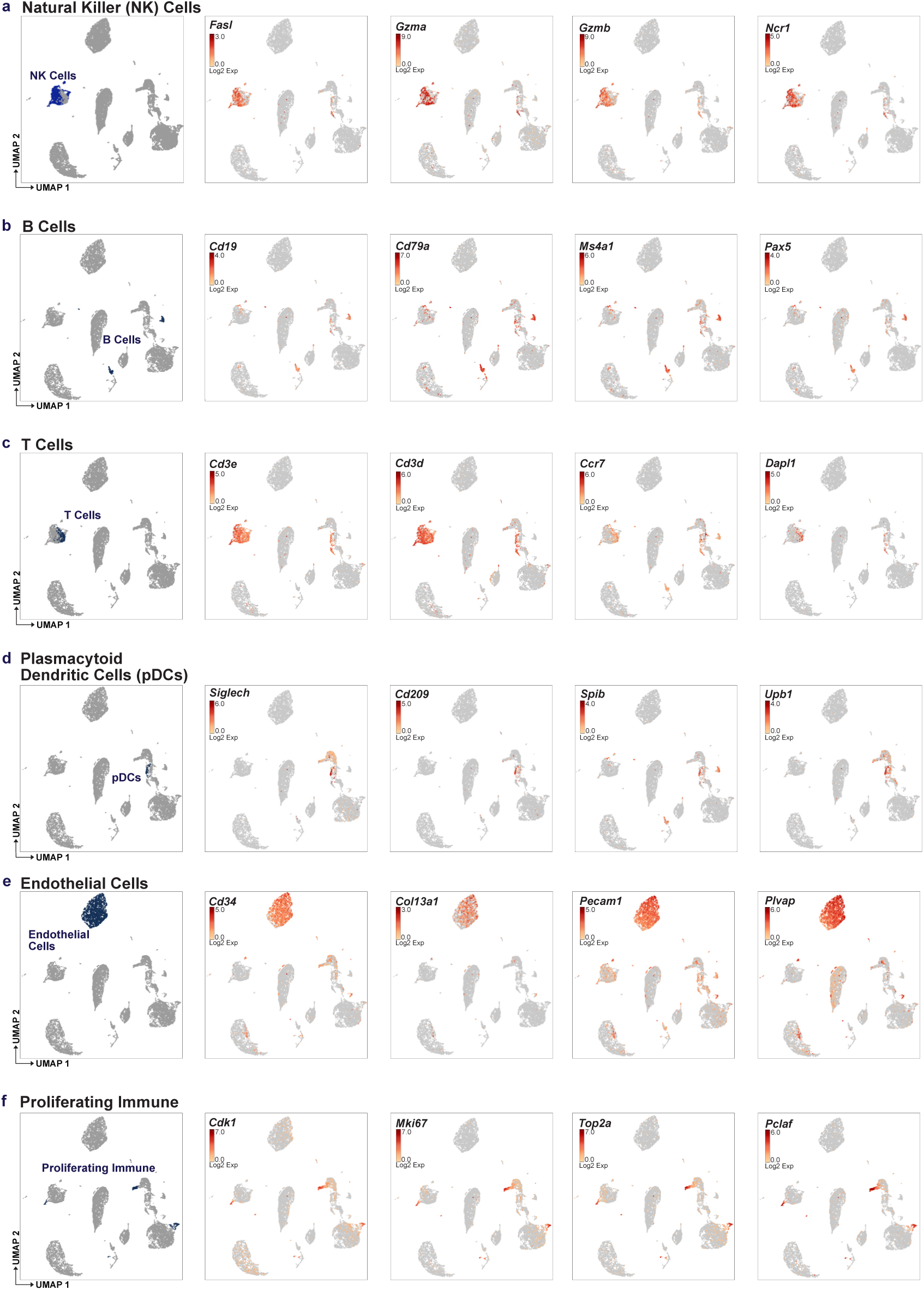
scRNAseq cluster validation of lymphoid, endothelial, and other lineages. UMAP represents major clusters as identified by established cell-type specific marker genes. (a) Natural killer (NK) cells identified by high expression of *Fasl*, *Gzma*, *Gzmb*, and *Ncr1*. (b) B Cells identified by high expression of *Cd19*, *Cd79a*, *Ms4a1*, and *Pax5*. (c) T cells identified by high expression of *Cd3e*, *Cd3d*, *Ccr7*, and *Dapl1*. (d) Plasmacytoid dendritic cells identified by high expression of *Siglech*, *Cd209*, *Spib*, and *Upb1*. (e) Endothelial cells identified by high expression of *Cd34*, *Col13a1*, *Pecam1*, and *Plvap*. (f) Proliferating immune identified by high expression of *Cdk1*, *Mki67*, *Top2a*, and *Pclaf*. Gene expression is represented as log2 expression as indicated in respective UMAPs.

**Extended Data Figure 7.**
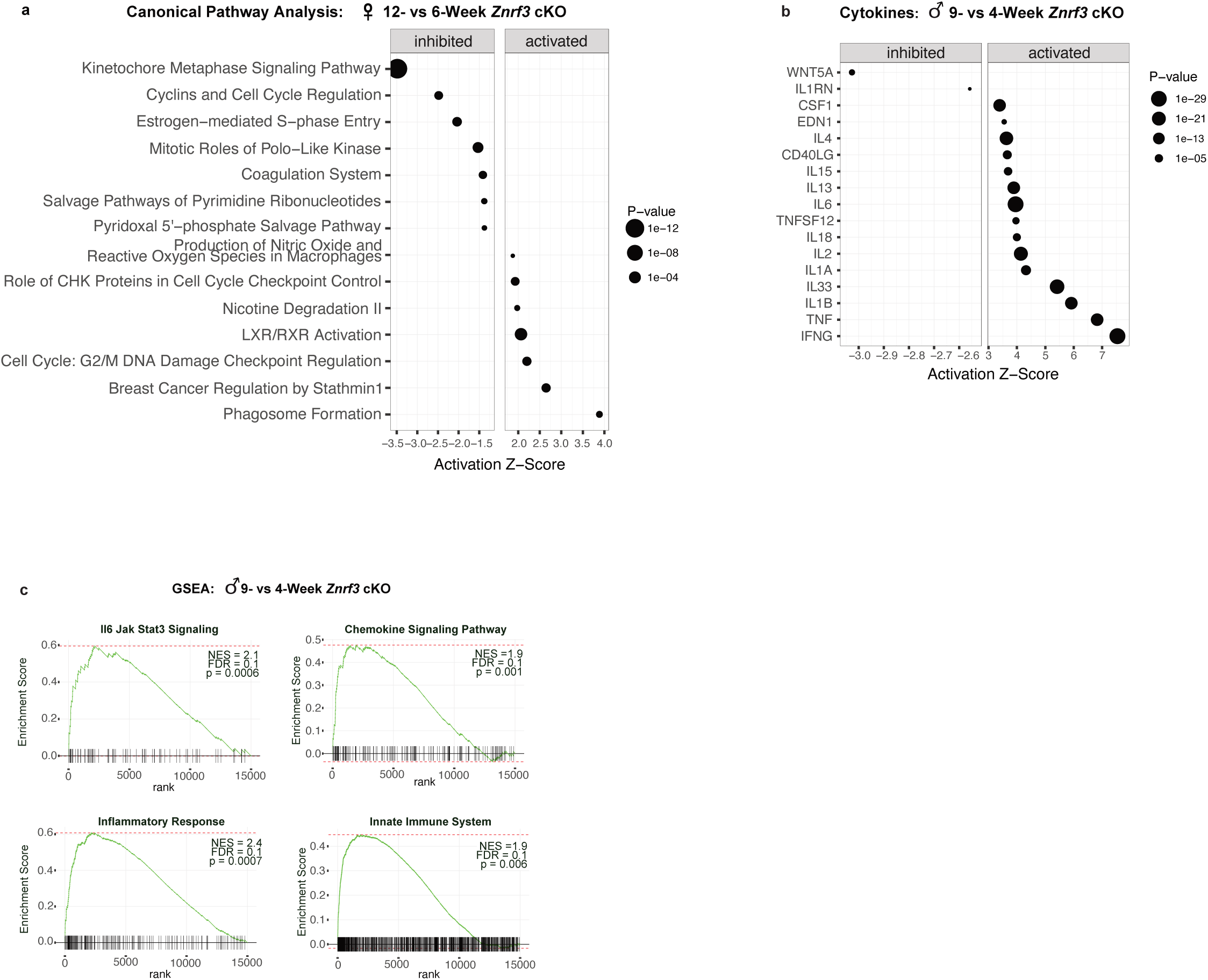
Inflammation and cytokine production in male senescent *Znrf3* cKO adrenal glands is further enhanced at 9-weeks. (a) IPA identifies the most significantly altered canonical pathways in 12- vs 6-week female *Znrf3* cKO adrenals. Similar pathways are altered earlier in male *Znrf3* cKOs, consistent with the more accelerated phenotype in males. (b) IPA identified the top activated and inhibited cytokines in 9- vs 4-week male *Znrf3* cKO adrenals. (c) GSEA for inflammatory signatures (IL6/JAK/STAT3 signaling, chemokine signaling, the inflammatory response, and the innate immune system) identifies positively enriched genes in 9- vs 4-week male *Znrf3* cKO mice. Data is representative of 4 biological replicates per group and statistical tests were performed using IPA, p<0.05.

**Extended Data Figure 8.**
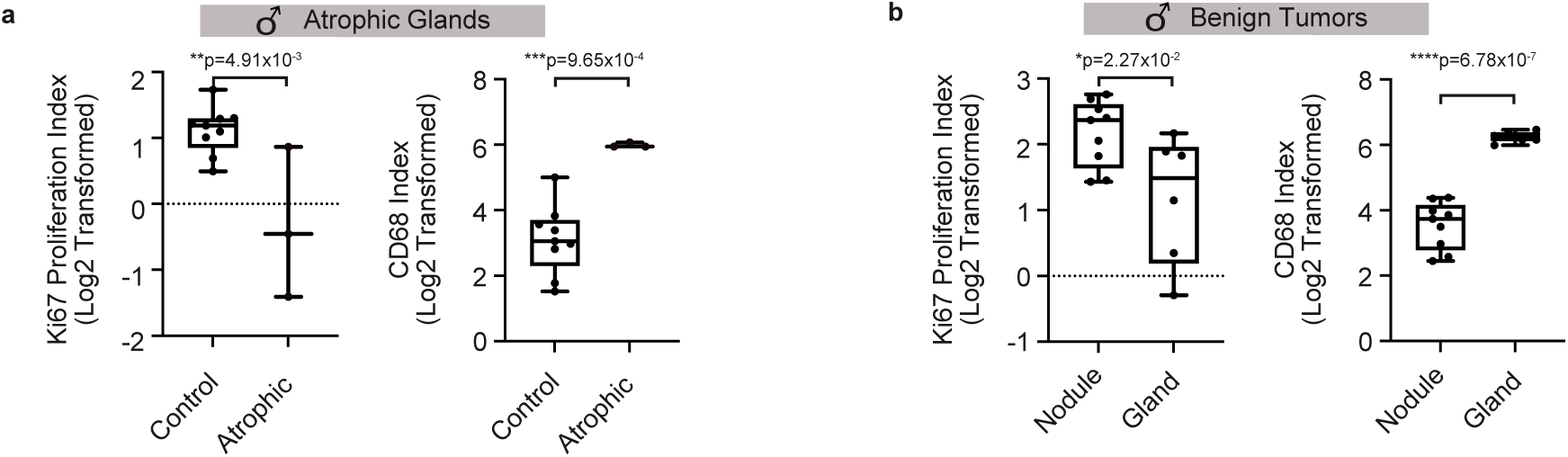
Proliferation and myeloid cell accumulation are inversely correlated in *Znrf3* cKO mice. (a) Atrophic *Znrf3* cKO adrenal glands have a significantly lower Ki67-index and higher CD68-index compared to age-matched controls. (b) In benign *Znrf3* cKO adrenals, nodules have a significantly higher Ki67-index and lower CD68-index compared to the background gland. Each dot represents an individual animal. Box and whisker plots represent mean with variance across quartiles. Statistical analysis was performed using two-tailed Student’s t-test.

**Extended Data Figure 9.**
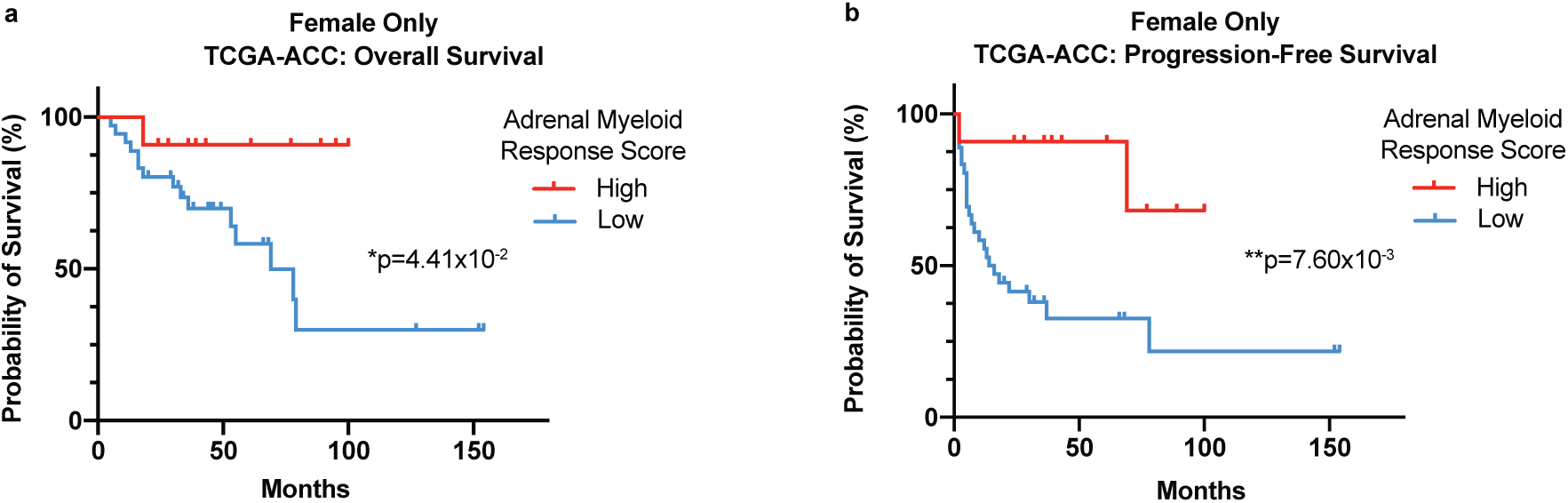
A low adrenal myeloid response score (AMRS) is associated with worse patient outcome in female ACC. Low adrenal MRS (AMRS) is associated with shorter (a) overall and (b) progression-free survival in female TCGA-ACC patients. Statistical analysis was performed using Log-rank Mantel-Cox test.

**Extended Data Figure 10.**
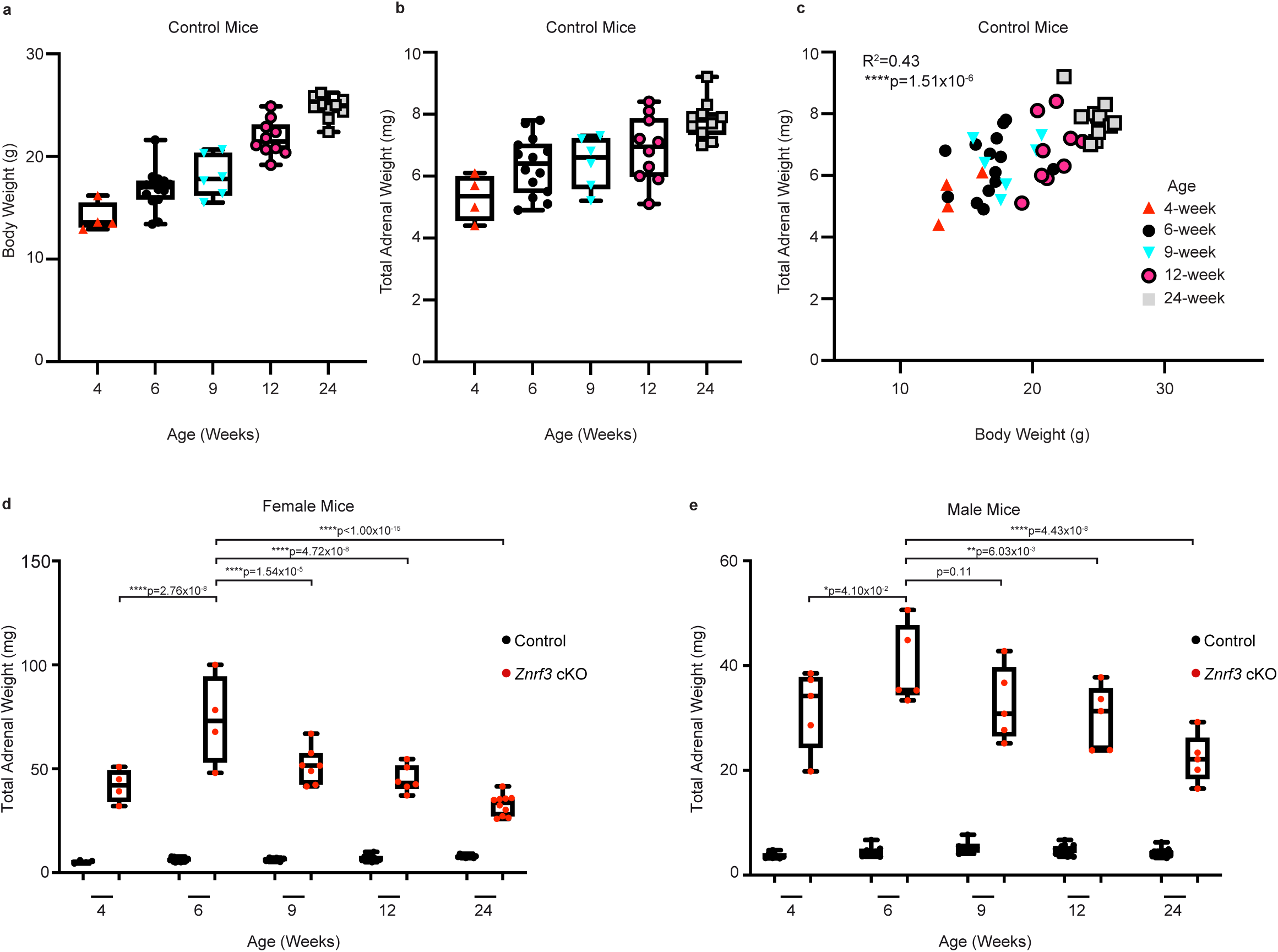
Extended adrenal weight analysis. (a) Body weight and (b) adrenal weight in normal, control mice progressively increase between 4- and 24-weeks of age, and (c) are significantly positively correlated. Raw total adrenal weight measurements in (d) female and (e) male cohorts from 4- to 24-weeks of age. Each dot represents an individual animal. Box and whisker plots represent mean with variance across quartiles. Statistical analysis was performed using simple linear regression (c) or two-way ANOVA followed by Tukey’s multiple comparison test (d-e).

